# Fexinidazole induced cytotoxicity is distinct from related anti-trypanosome nitroaromatic drugs

**DOI:** 10.1101/2023.10.09.561529

**Authors:** Indea Rogers, Kenna Berg, Hayley Ramirez, G A Hovel-Miner

## Abstract

Nitroaromatic drugs are of critical importance for the treatment of trypanosome infections in Africa and the Americas. Fexinidazole recently joined benznidazole and nifurtimox in this family when it was approved as the first oral therapy against Human African trypanosomiasis (HAT). Nitroaromatic prodrugs are bioactivated by the trypanosome-specific type I nitroreductase (NTR) enzyme that renders the compounds trypanocidal. A caveat to the specificity of NTR activation is the potential for drug resistance and cross-resistance that can arise if NTR expression or functionality is altered through mutation. The outcomes of NTR bioactivation of nitroaromatic compounds is variable but can include the formation highly reactive open chain nitriles that can damage biomolecules including DNA. A proposed mechanism of action of nitroaromatic compounds is the formation of reactive oxygen species (ROS) resulting in the formation of trypanocidal levels of DNA damage. Fexinidazole made its way to clinical approval without a significant interrogation of its effects on trypanosome biology and a limited understanding of its mechanism of action. Early reports mentioned fexinidazole potentially affects DNA synthesis but without supporting data. In this study, we evaluated and compared the cytotoxic effects of nifurtimox, benznidazole, and fexinidazole on *Trypanosoma brucei* using in vitro analyses. Specifically, we sought to differentiate between the proposed effects of nitroaromatics on DNA damage and DNA synthesis. Toward this goal we generated a novel γH2A-based flow cytometry assay that reports DNA damage formation in conjunction with cell cycle progression. Here we report that fexinidazole’s cytotoxic outcomes are distinct from the related drugs nifurtimox and benznidazole. Specifically, we show that fexinidazole treatment results in a pronounced defect in DNA synthesis that reduces the population of parasites in S phase. In contrast, treatment with nifurtimox and benznidazole appear accumulate DNA damage early in cell cycle and result in a defective G_2_ population. The findings presented here bring us closer to understanding the anti-trypanosomatid mechanisms of action of nitroaromatic compounds, which will promote improved drug design and help combat potential drug resistance in the future. Our findings also highlight DNA synthesis inhibition as a powerful anti-parasitic drug target.

## 1. INTRODUCTION

Fexinidazole has recently joined nifurtimox and benznidazole in a group of nitroaromatic compounds critical to the treatment of human trypanosome infections (Deeks, 2019a). Trypanosomes are unicellular protists in the group kinetoplastida that cause the significant human diseases: African trypanosomiasis (*Trypanosoma brucei spp.*), American trypanosomiasis (Chagas disease, *Trypanosoma cruzi*), and Leishmaniases (*Leishmania spp.*). These related human parasites differ in global distribution and diseases progression, yet they are all transmitted by insect vectors and have an initial acute phase followed by a chronic infection. Human African Trypanosomiasis (HAT) is unique in that parasite growth and replication is entirely extracellular and HAT infections are invariably fatal to the host in the absence of drug treatment. The majority of *T. cruzi* and *Leishmania* human infections are self-resolving in the acute phase and progress to chronic, life-threatening infections, in about 25-30% of cases (Rycker et al., 2023). HAT infections are caused by the two subspecies *T. b. gambiense* (g-HAT) and *T. b. rhodesiense (*r-HAT). HAT begins as a bloodstream infection (stage I) and progress to an infection of the central nervous system (stage II), which results in a parasite infection in the brain, and ultimately, death. *T. b. gambiense* is a far more prevalent cause of HAT and results in chronic infections lasting months or even years, whereas r-HAT accounts for fewer cases but can rapidly progress often causing death within weeks.

The efficacy of nitroaromatic compounds against trypanosome infections dates to the 1950s with benznidazole used to treat Chagas diseases (*T. cruzi*) for over 40 years. Nifurtimox has been used in the treatment of Chagas and HAT (Alirol et al., 2013; Patterson and Wyllie, 2014a), more recently, in the form of nifurtimox and eflornithine combination therapy (NECT) (Alirol et al., 2013; Babokhov et al., 2013). In 2019, fexinidazole was approved as the first oral monotherapy against g-HAT with much enthusiasm over the new therapies’ potential role in eradicating g-HAT, a long-standing WHO initiative (Deeks, 2019b; Deeks and Lyseng-Williamson, 2019). However, fexinidazole’s cytotoxic effects have not been established, its mechanism of action (MoA) is unknown, and the potential for drug resistance remains a concern.

Fexinidazole is a 5-nitroimidazole whose potency against trypanosomatids was demonstrated in vivo and in vitro in 1983 (Raether and Seidenath, 1983). The compound was not pursued clinically at that time because of broad concerns over the potential host toxicity of nitroaromatic compounds. Nitro groups can be highly reactive possessing polarity, hydrogen bonding, and electron-withdrawing chemical properties (Whitmore and Varghese, 1986). Concerns about potential host toxicity of anti-trypanosome nitro compounds were later mitigated by the discovery that trypanosomatids harbor a bacteria-like type I nitroreductase (NTR), which is required for the bioactivation of nifurtimox and benznidazole in a manner that is distinct from the mammalian type II NTR (Wilkinson et al., 2008). Thus, NTR activation of nitroaromatic prodrugs in trypanosomes occurs through sequential two-electron reduction of nitro groups to produce amines via nitroso and hydroxylamine intermediates (Patterson and Wyllie, 2014b). Incubation of trypanosome NTR with nifurtimox results in the formation of reactive open-chain nitriles (Hall et al., 2011). Whereas benznidazole bioactivation by trypanosome NTR forms a dihydro-dihydroxyl imidazole which exists in equilibrium with glyoxal and guanidine products, where glyoxal is highly toxic and capable of modifying biomolecules including nucleotides (Hall and Wilkinson, 2011). The trypanocidal consequences of nifurtimox and benznidazole bioactivation are predicted to arise from the formation reactive oxygen species (ROS) resulting in toxic levels of DNA damage (Hall et al., 2011). While fexinidazole is expected to undergo trypanosome specific NTR activation, the resulting chemical outcomes have not yet been reported.

Due to an urgent need for a new treatment for second stage HAT, to replace the highly toxic drug melarsoprol, fexinidazole followed an unusual path to clinical usage. In a collaboration between Sanofi and the Drugs for Neglected Diseases initiative (*DNDi*), fexinidazole was proven safe and efficacious for the oral treatment of g-HAT within less than 10 years of clinical trials (Deeks, 2019b). Because of this pathway to approval, little basic research was published on fexinidazole resulting in a considerable knowledge gap regarding the drug’s mechanism of action and trypanocidal effects. What is known is that resistance to fexinidazole can arise rapidly in vitro and resistant clones can display cross-resistance against other nitroaromatic compounds, nifurtimox and benznidazole specifically, which may or may not arise via NTR mutations (Sokolova et al., 2010a; Wyllie et al., 2016a). Thus, it remains unclear precisely how any of the clinically relevant anti-trypanosomatid nitro drugs kill trypanosomes.

Here we evaluate the trypanocidal outcomes of fexinidazole treatment in comparison with nifurtimox and benznidazole. Specifically, in vitro assays of the model African trypanosome *T. b. brucei* were interrogated to determine the effects of nitroaromatic drugs on cell cycle, DNA synthesis, and DNA break formation; using a novel γH2A-based flow cytometry assay developed herein. Drug concentrations that eliminate parasites after 4-6 days of treatment were used to evaluate drug-induced cytotoxicity phenotypes within 24 and 48 hours. Our findings indicate that, while nifurtimox and benznidazole have similar cytotoxic outcomes, fexinidazole’s trypanocidal mechanism is distinct. These discoveries are impactful because they bring us closer to understanding the MoA of all 3 drugs and provide insights into chemical improvements in future anti-trypanosome drug development that could be designed based on the nitroaromatic compound scaffold.

## 2. RESULTS

### Fexinidazole treatment results in spontaneous resistance and cross-resistance

Fexinidazole and the related nitroaromatics benznidazole and nifurtimox are proposed to kill trypanosomatid parasites by way of ROS induced DNA damage (Zuma et al., 2019). To begin investigating this prediction we sought to establish treatment conditions in each drug that killed *T. brucei* parasites in vitro in a similar time frame. Cell viability assays using AlamarBlue reported EC_50_ values of 7 μM, 35 μM, and 3 μM for nifurtimox, benznidazole, and fexinidazole, respectively (Supplemental Figure 1). Thus, we tested a range of drug concentrations, starting at approximately 5x the EC_50_, to identify conditions that resulted in complete parasite death between 3-5 days of treatment. Initial concentrations were determined using cumulative growth assays over 5 days to identify high, mid, and low drug concentrations (FIG. 1A – Low in green, Mid in blue, and High in Red). In most instances the low concentration did not eliminate all the parasites within 1 week. Whereas most high concentrations eliminated all countable parasites between 48 and 72 hours. These concentrations were then used to evaluate death in continuous culture for 10-12 days, where the media and drug were replaced completely every 3 days (FIG. 1B). The high and mid drug treatment concentrations appeared to kill all parasites in all drugs by day 5 or 6 (FIG. 1B). However, both fexinidazole and nifurtimox resulted in the formation of resistant survivor populations at around day 6 or 7 that ultimately were able to grow robustly in each drug, respectively (FIG. 1B). Benznidazole did not result in drug resistance under these conditions. Fexinidazole resistant survivors from low (20 μM) and mid (50 μM) drug concentrations were isolated and tested for their ability to cause cross-resistance to nifurtimox and benznidazole. Both populations of fexinidazole drug resistant survivors resulted in cross-resistance to nifurtimox and benznidazole, with some variations in the timing of death (FIG. 1C – orange and purple lines). Because all nitroaromatic drugs are activated in trypanosomes by a type I NTR enzyme, and this has been shown to cause drug resistance, we PCR amplified the NTR enzyme encoding gene from the fexinidazole resistant isolates and sequenced the NTR gene to determine if it harbored mutations that could account for the observed drug resistance and cross resistance. We did not find any mutations in the NTR gene that would be indicative of the observed resistance phenotypes (data not shown). However, our analysis did not exclude possible mutations affecting NTR expression.

**FIGURE 1.**
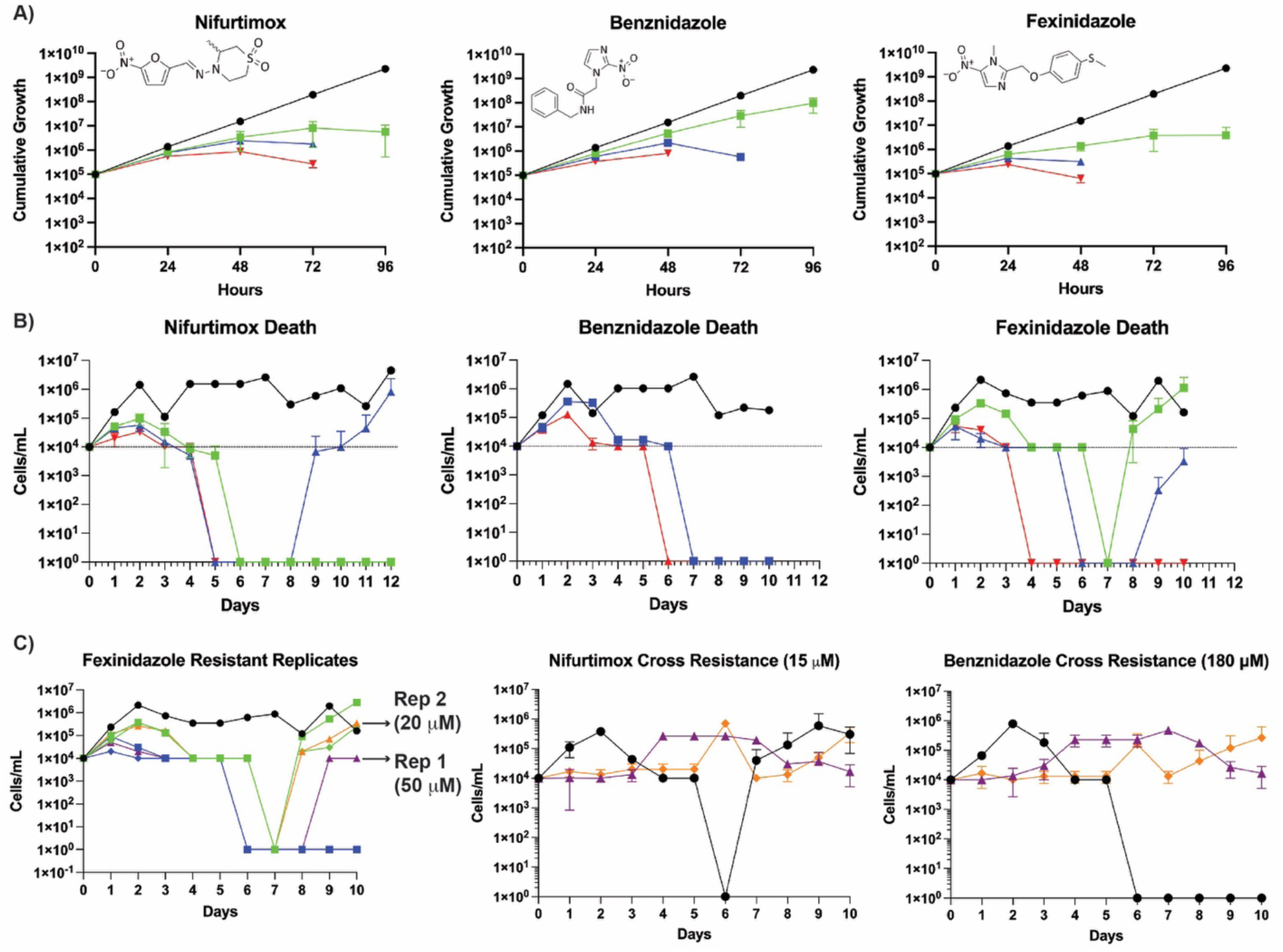
*T. brucei* killing in nitroaromatic drugs and fexinidazole cross resistance. A) Cumulative Growth Assays were conducted during treatment with nifurtimox (6 μM green, 9 μM blue, and 12 μM red), benznidazole (60 μM green, 120 μM blue, and 180 μM red), and fexinidazole (20 μM green, 50 μM blue, and 70 μM red) daily dilution of parasites to 100,000 cells/mL and quantification of their total growth over 5 days. The termination of a line before 96 hours indicates parasites that were too few to pass. B) *T. brucei* death was monitored for 10-12 days during treatment with nifurtimox (9 μM green, 12 μM blue, and 15 μM red), benznidazole (120 μM blue and 180 μM red), or fexinidazole (20 μM green, 50 μM blue, and 70 μM red). Every three days parasites were replenished with fresh media and drug. C) Fexinidazole drug resistant parasites were isolated from 20 μM (purple) and 50 μM (orange) and grown in the presence of nifurtimox (15 μM) or benznidazole (180 μM) in comparison to parental control parasites (black).

Thus, we empirically determined drug treatment conditions for nifurtimox, benznidazole, and fexinidazole that eliminate *T. brucei* parasites in vitro between 3-5 days of treatment. As a point of comparison, we established the treatment conditions for eflornithine whose molecular target is known to be ornithine decarboxylase (Krauth-Siegel and Comini, 2008)and distinct from the nitroaromatic drugs (Supplemental Figure 2). The drug treatment conditions established here will be utilized throughout this study to evaluate drug-induced cytotoxicity.

### Fexinidazole treatment decreases the S phase population

To better understand the cytotoxic effects of fexinidazole on *T. brucei* we evaluated the cell cycle effects of 3 drug concentrations, 20 μM, 50 μM, and 70 μM that eliminate parasites after 7, 6, and 4 days of treatment, respectively. These were compared with Untreated Control (UT) (FIG. 2 – bottom right) whose populations consisted of 4.9% AN, 51.7% G_1_, 15.6% S phase, and 33.4% G_2_ cells. Following 24 hours of fexinidazole treatment a minor reduction in the population of G_2_ cells was observed (FIG. 2A). However, by 48 hours of drug treatment the percent of cells in S phase was reduced by approximately 5%, from 15% in UT to 10% in fexinidazole treated parasites (FIG. 2). Overlay of cell cycle traces from UT and fexinidazole treated cells illustrates that loss of S phase cells is the primary defect across all drug concentrations by 48 hours post-treatment (FIG. 2B).

**FIGURE 2.**
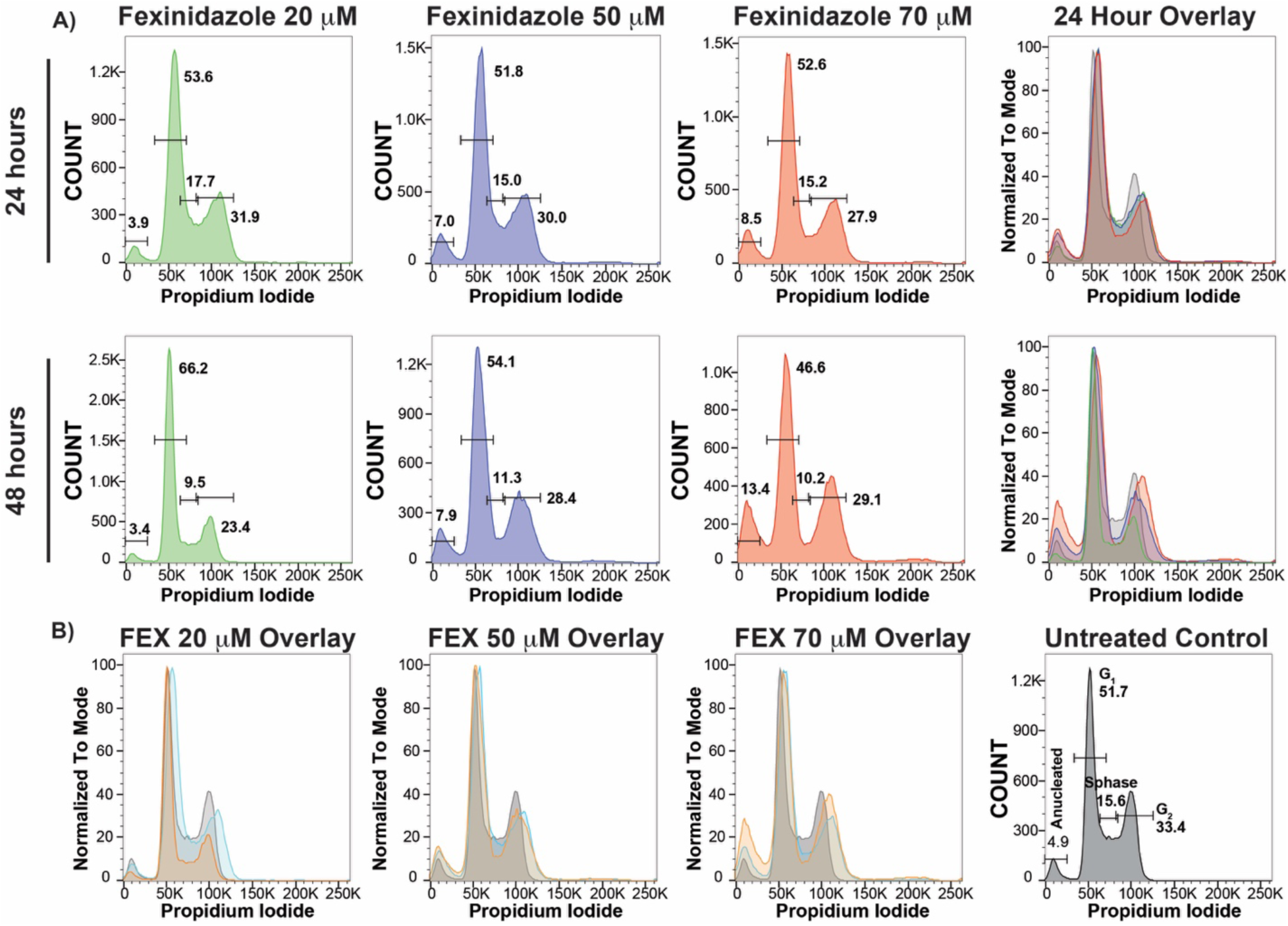
Fexinidazole cell cycle analysis after 24 and 48 hours of drug treatment. Cell cycle analysis based on propidium iodide DNA content staining evaluated by flow cytometry for untreated control (UT), with cell cycle populations shown as percent of total and (A) Fexinidazole concentrations (20 μM green, 50 μM blue, and 70 μM red) analyzed at 24 hours and 48 hours. (B) Comparisons between concentrations and time points are also shown.

### Nifurtimox and benznidazole treatment decreases cells in G_2_

To investigate the proposed MoA of nifurtimox and benznidazole, we first interrogated their cell cycle phenotypes at mid and high drug concentrations after 24 and 48 hours of treatment. Within 24 hours of treatment both nifurtimox and benznidazole displayed a decrease in their G_2_ cell population from approx. 33% of UT (FIG. 2) to approx. 24% in nifurtimox and 28% in benznidazole (FIG. 3A & 3B). The drug induced defect in G_2_ cell populations was more pronounced after 48 hours of treatment with nifurtimox at approx. 18% and benznidazole having approx. 22% of cells in G_2_ (FIG. 3 – 48 hour). In comparison with UT cells, which have approx. 33% of their cells in G_2_, this is a reduction of over 20% for nifurtimox and 10% for benznidazole. The observed reduction in G_2_ populations during these drug treatments was correlated with an increase in cells in G_1_ and S phase (FIG. 3).

**FIGURE 3.**
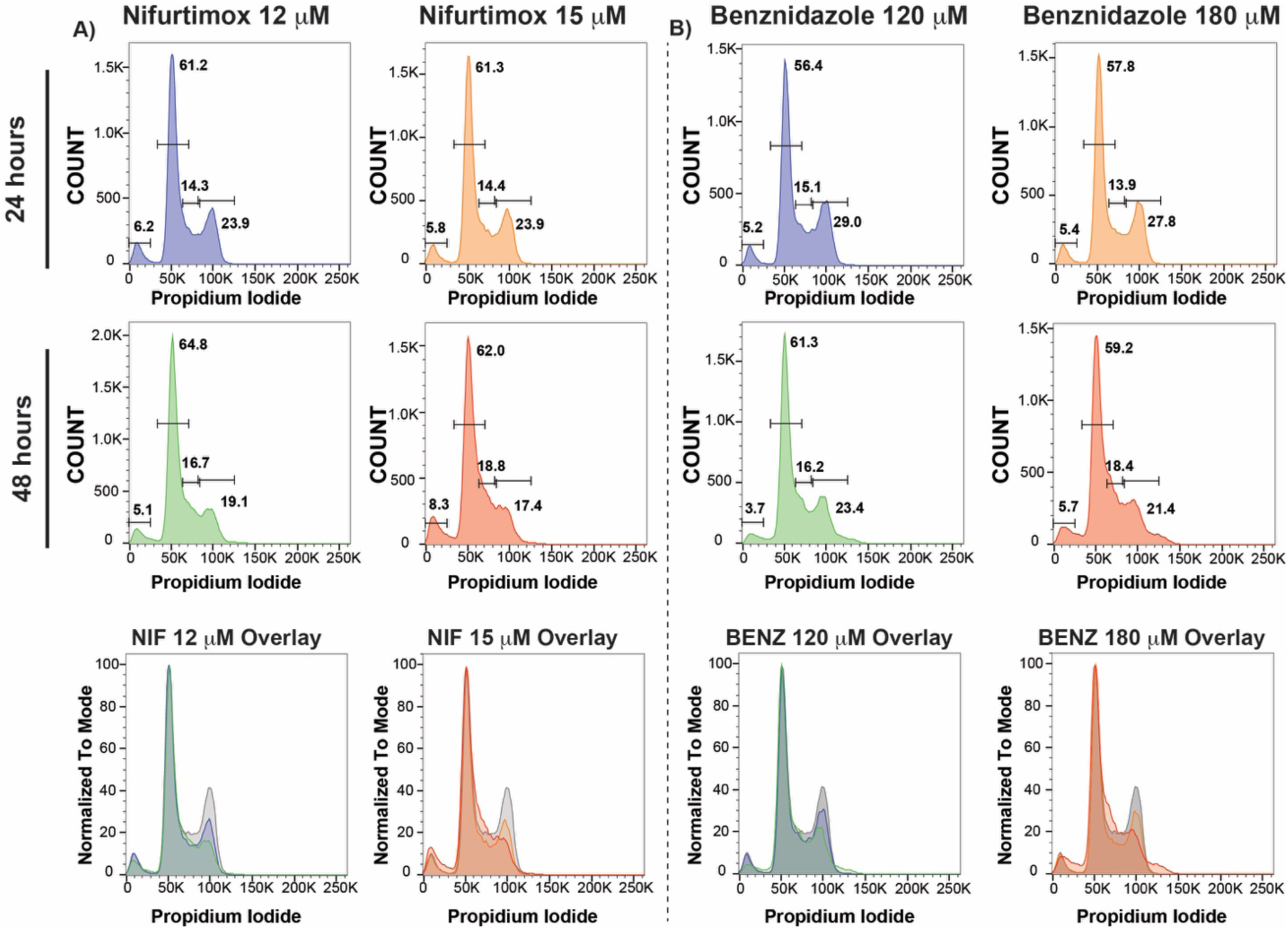
Effects of nifurtimox and benznidazole treatments on cell cycle progression. A) Analysis of cell cycle effects of nifurtimox at 12 μM after 24 hours (blue) and 48 hours (green) or nifurtimox at 15 μM after 24 hours (green) or 48 hours (red). B) Cell cycle analysis of benznidazole at 150 μM after 24 hours (blue) and 48 hours (green) or nifurtimox at 180 μM after 24 hours (green) or 48 hours (red). Bottom - Overlays of nifurtimox (NIF) or benznidazole (BENZ).

Overlay of cell cycle histograms of benznidazole and nifurtimox cell cycle analysis demonstrates the progressive loss of G_2_ cells over increasing concentration and duration of treatment (FIG. 3 – bottom panels). While the drug induced cell cycle phenotype of nifurtimox is slightly more pronounced, the pattern for both drugs is highly similar and indicates a cessation of growth arising from a cell cycle stall in S phase. As a point of comparison, this was not the case for during eflornithine treatment, which results in a large accumulation of parasites in G_2_ (Supplemental Figure 2), demonstrating that not all anti-trypanosome cytotoxicity proceeds with the same cell cycle kinetics. The loss of G_2_ cells may support the prediction that nitroaromatic-induced DNA damage results in trypanosomatid death for nifurtimox and benznidazole.

### Fexinidazole’s effects on cell cycle are distinct from related nitroaromatics

To further investigate the cell cycle defects associated with each of the 3 clinically relevant anti-trypanosomatid nitroaromatic drugs, we sought to directly compare each drug and evaluate the statistical significance of our observations (FIG. 4). The cell cycle profiles of nitroaromatic drugs were compared at 24 and 48 hours. We observed that while nifurtimox, benznidazole, and fexinidazole appear somewhat similar after 24 hours, with minor reductions in both S phase and G_2_ populations, by 48 hours of treatment fexinidazole results in a distinct cell cycle profile (FIG. 4A). Specifically, fexinidazole results a large, and perhaps broader, G_2_ population and a marked decrease in the S phase population. By in large, these observations were statistically significant when compared among at least 3 biological replicates. For comparison we also included eflornithine treatment at 100 μM. Only fexinidazole (70 μM, 48 hrs) and eflornithine resulted in a statistically significant increase on the formation of AN cells (FIG. 4B). The most significant comparisons among nitroaromatic drugs were on the S phase and G_2_ populations. Both nifurtimox and benznidazole demonstrated a statistically significant and reproducible reduction in G_2_ populations at 48 hours in both drug concentrations tested (FIG. 4B). Whereas fexinidazole’s G_2_ defect was only significant in the lowest concentration following 24 hours of treatment. The most pronounced cell cycle phenotype associated with fexinidazole treatment was a statistically significant reduction in S phase populations by approx. 20% compared with UT cells (FIG. 4B). Thus, while nifurtimox and benznidazole result in similar cell cycle defects, the 5-nitroimidazole fexinidazole is unique among the nitroaromatics in its reduction of the S phase population.

**FIGURE 4.**
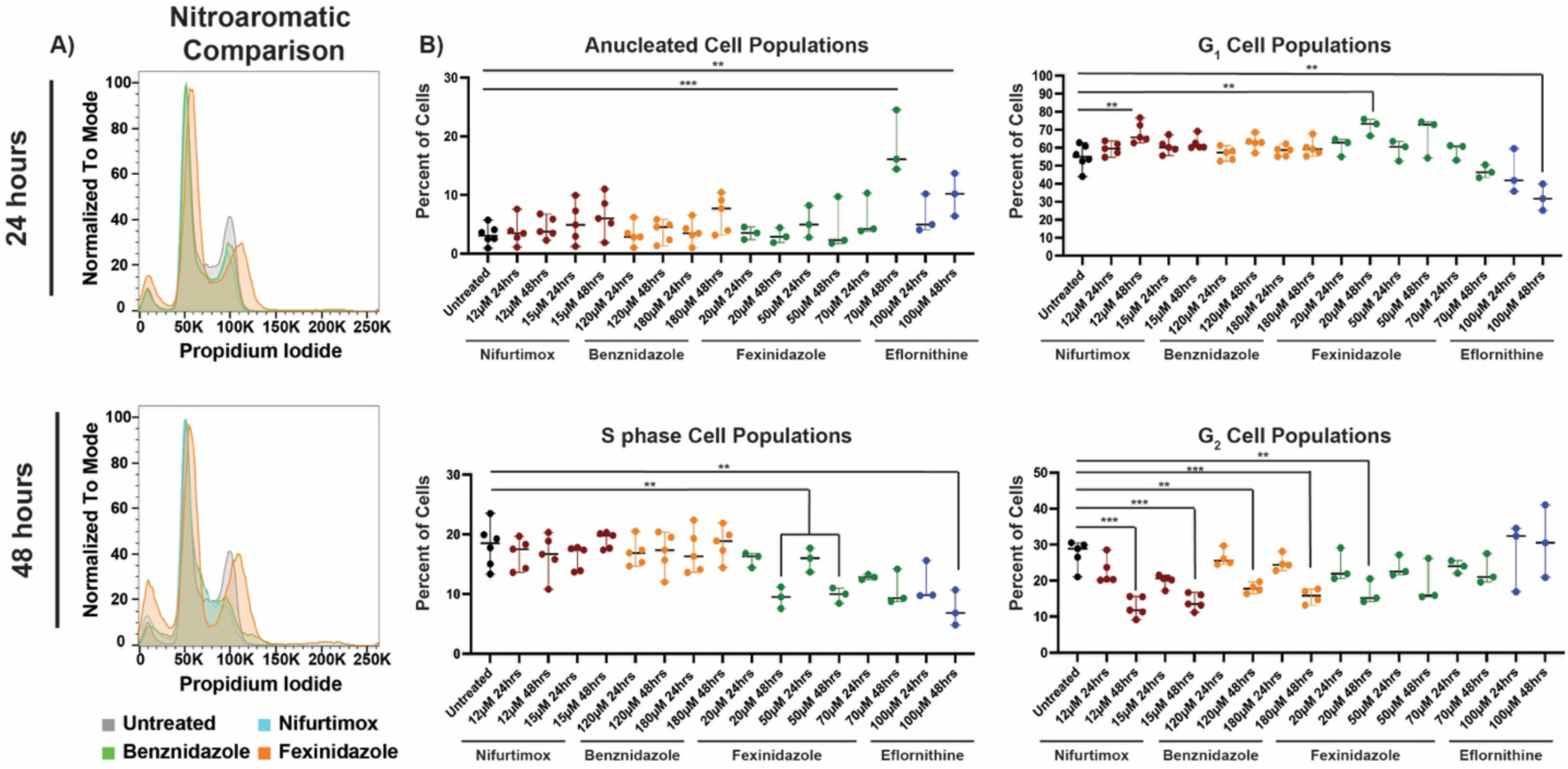
Comparative analysis and statistical significance of cell cycle effects. A) Overlays of Untreated (grey), nifurtimox (blue, 15 μM), benznidazole (green, 180 μM), and fexinidazole (orange, 70 μM) at 24 and 48 hours of treatment. B) Statistical comparison of proportions of cell cycle populations (from flow cytometry analysis) among biological replicates of drug concentrations analyzed. Dots represent a single replicate of each untreated (black) or drug treatment conditions: nifurtimox (red), benznidazole (orange), fexinidazole (green), and eflornithine (blue). Error bars show the mean and standard deviation. Statistical significance calculated based on students t-test for comparison represented by line or brackets with pval<0.001 (***) or pval<0.01 (**) indicated.

### Fexinidazole treatment results in changes in cell cycle progression by IF

To evaluate the cell cycle phenotypes of nitroaromatic drugs in greater detail, we analyzed the cellular composition of kinetoplasts and nuclei by immunofluorescence (IF) microscopy. Trypanosomatids are unique among protists in that they harbor a mitochondrial DNA containing sub-organelle called the kinetoplast, which defines their phylogeny. During the cell cycle, S phase begins with the elongation and subsequent division of the kinetoplast prior to the completion of nuclear DNA synthesis and division of nuclei. Thus, the trypanosome cell cycle can be monitored based on the number of kinetoplasts and nuclei per cell. Trypanosomes in G_1_ harbor one kinetoplast and one nucleus (1K1N), entry into S phase is characterized by cells harboring 2K and 1N (2K1N), and completion of S phase (G_2_) results in parasites with 2K and 2N (2K2N), after which parasites normally proceed to cytokinesis (Hammarton, 2007; Jones et al., 2014). To evaluate cell cycle progression by IF microscopy, cells were stained with an anti-VSG-2 coat antibody, to visualize the cell surface, and DAPI to visualize both the kinetoplasts and nuclei (FIG. 5). At least 100 cells were analyzed from each drug treatment condition, for which the number of kinetoplasts and nuclei counted for each cell. In addition, during drug treatments, there are often increases in “other” cell populations, which can include AN, multinucleated, and additional forms of aberrant drug induced cytotoxicity (Examples can be seen in Supplemental Figure 3). Analysis of IF microscopy for UT parasites resulted in 58% 1K1N (G_1_), 27% 2K1N (S phase), 12% 2K2N (G_2_), and 2% other (Supplemental Figure 3). Both nifurtimox and benznidazole displayed a progressive loss of their 2K2N population upon increasing concentration and duration of treatment down to 8% for nifurtimox and 6.5% for benznidazole. The decreased 2K2N populations during these drug treatments supports the loss of G_2_ observed by cell cycle flow cytometry analysis. In contrast, fexinidazole treatment resulted in an increase in 2K2N harboring cells to 20-25%, in conjunction with a significant reduction in 2K1N parasites from 27% in untreated down to 7-9% in fexinidazole treated cells (FIG. 5B). The observed decrease in 2K1N cells during fexinidazole treatment correlates with the decrease in the S phase population visualized by flow cytometry. How the fexinidazole induces a decrease in cells in S phase (FIG. 2, 4, & 5) and results in an increase in 2K2N harboring cells (FIG. 5B), with an apparent cell cycle stall, is the subject of further consideration.

**FIGURE 5.**
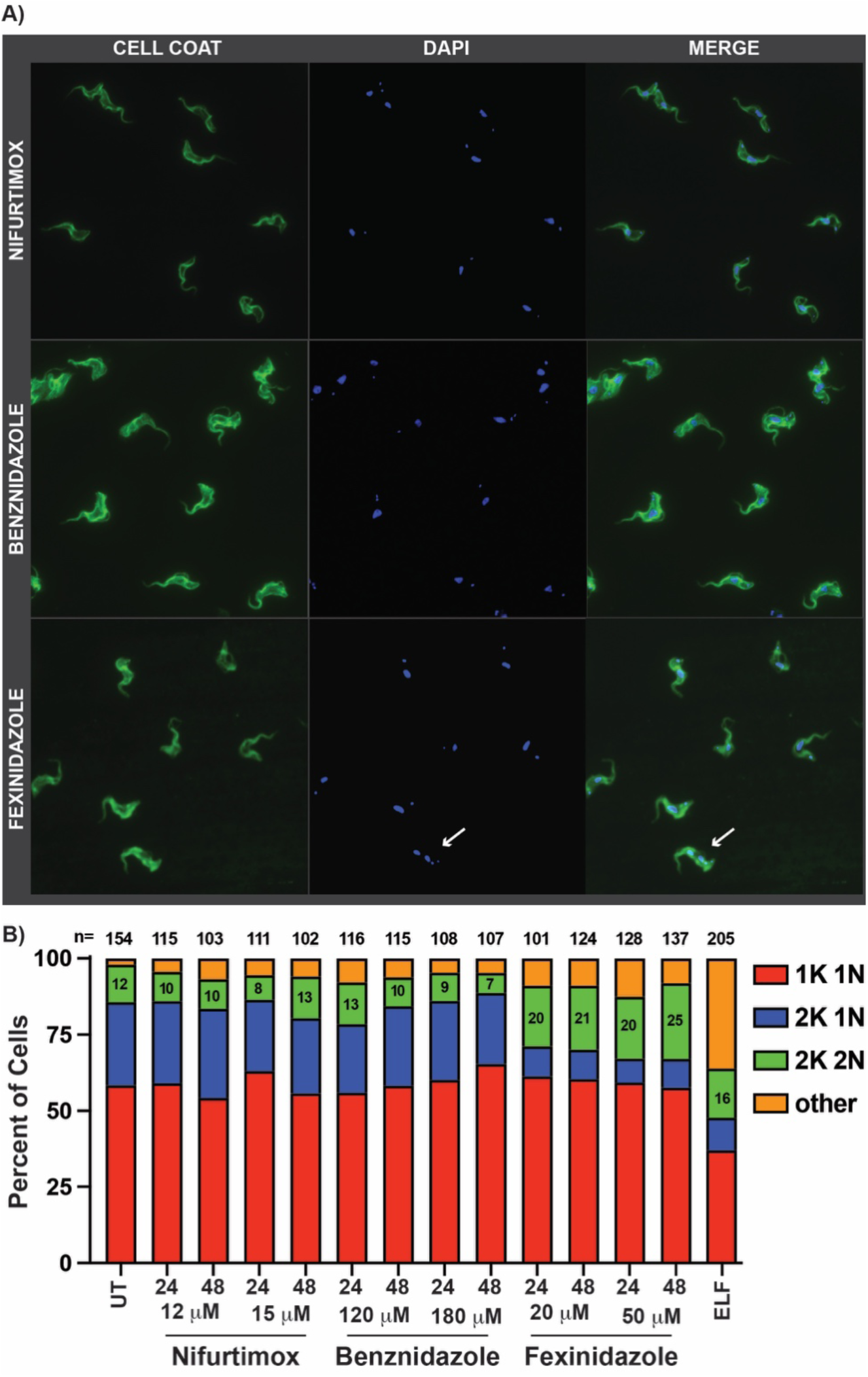
Nitroaromatic drugs have different effects on parasite kinetoplast (K) and nuclei (N) composition. A) Sample IF microscopy images for *T. brucei* treated with nifurtimox (15 μM), benznidazole (180 μM), or fexinidazole (50 μM). For UT see IF microscopy Supplemental Figure 3. B) Graphical analysis of kinetoplast (K) and nuclei (N) counts per cell analyzed in FIJI with 1K1N (red), 2K1N (blue), 2K2N (green), and patters outside these parameters shown as “other” (orange). At least 100 parasites were counted per condition with the total counted parasites shown as n= value at the top each bar on the graph. The percent of the population harboring 2K2N are shown in the green section of the bar graph.

### Fexinidazole treatment causes a specific DNA synthesis defect

In a previous study, we discovered that melarsoprol, a classic arsenical drug critical in the treatment of second stage *T. b. rhodesiense*, results in the rapid inhibition of DNA synthesis (Larson et al., 2021). Melarsoprol’s inhibition of DNA synthesis was dependent on the trypanothione pathway and could be partially alleviated by the overexpression of γ-glutamylcysteine synthetase, which increases the levels of available trypanothione (Shahi et al., 2002). At that time, we reported a partial reduction in DNA synthesis during fexinidazole treatment and that the effect did not appear to be associated with trypanothione (Larson et al., 2021). Based on cell cycle observations herein, that fexinidazole decreases S phase cell populations, we sought to evaluate the effects of all three anti-trypanosome nitroaromatic drugs on DNA synthesis. We employed a flow cytometry-based bromodeoxyuridine (BrdU) incorporation assay, in which parasites are pulsed with the alternative base BrdU and then DNA synthesis is measured using an anti-BrdU antibody to determine the proportion of BrdU positive cells (Silva et al., 2017; Larson et al., 2021). Cells are also stained with DAPI to identify the specific populations of S phase cells with BrdU incorporated. Analyses were based on the formation of an anti-BrdU positive second peak on a histogram (FIG. 6B) and the S phase specific BrdU staining population when graphed against DAPI (FIG. 6C). Based on these metrics we observe that UT parasites grown for 24 hours consist of between 33% (DAPI vs. anti-BrdU) to 36% (anti-BrdU histogram) DNA synthesizing cells. We then evaluated the proportion of DNA synthesizing cells following 24 hours of treatment with nifurtimox (15 μM), benznidazole (180 μM), and fexinidazole (50 μM) (FIG. 6). Treatment with benznidazole did not result in any loss of the DNA synthesizing parasite population (FIG. 6), whereas treatment with nifurtimox resulted in a minor reduction to 17-21% of parasites undergoing DNA synthesis. In contrast, 24 hours of fexinidazole (50 μM) treatment resulted in a near complete loss of DNA synthesizing cells down to less than 5% of the total cell population (FIG. 6B & 6C). Overlay of anti-BrdU positive populations from UT and drug treated parasites clearly show that, while the shape of the populations may differ for nifurtimox and benznidazole, fexinidazole is unique in its nearly complete loss of DNA synthesizing cells (FIG. 6E & 6F). These data support the cell cycle data that S phase cell populations are decreased during fexinidazole treatment and raise questions regarding how the drug induces DNA synthesis inhibition while resulting in an accumulation of apparently cytokinesis stalled parasites harboring 2K2N.

**FIGURE 6.**
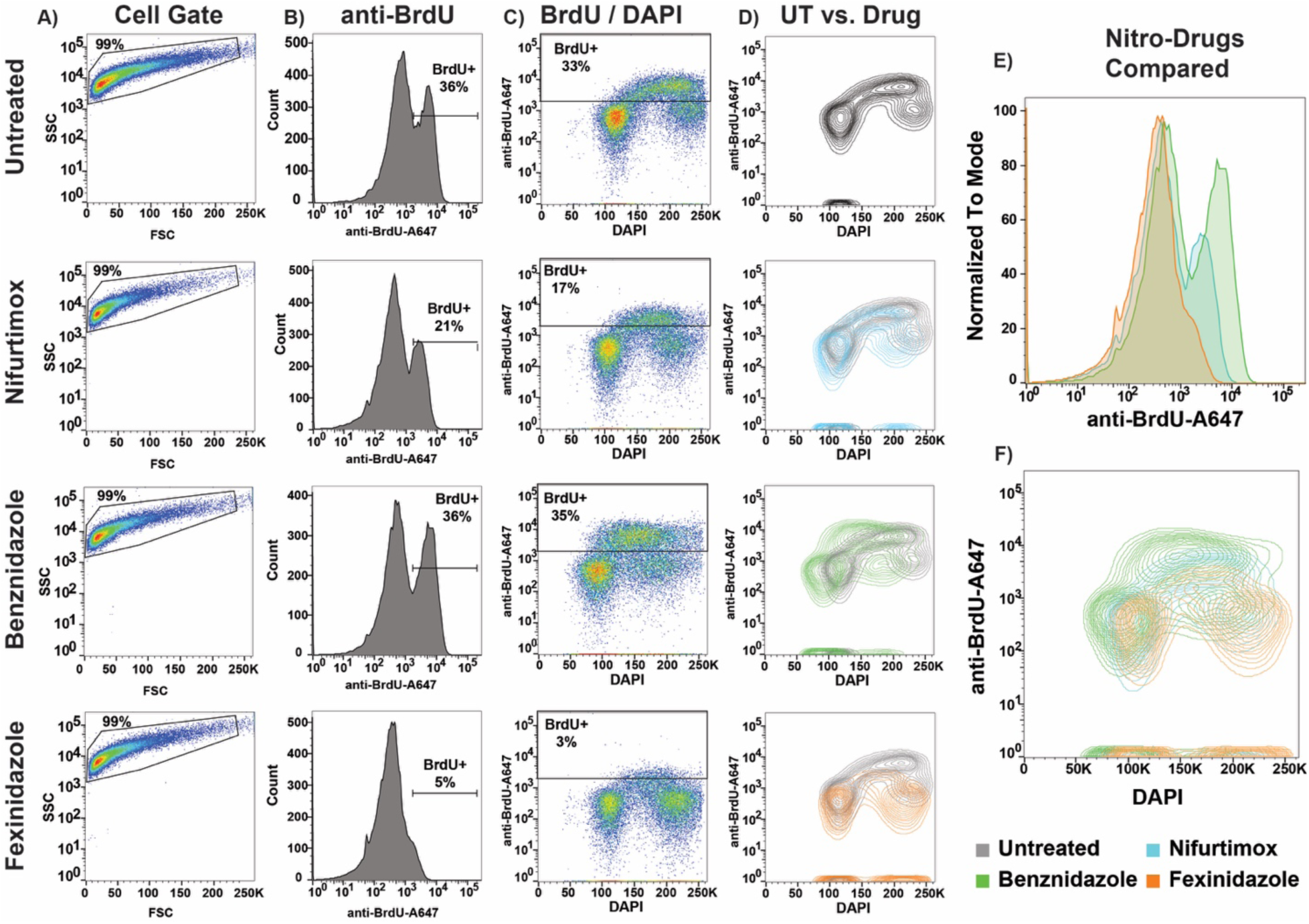
Analysis of DNA synthesis during nitroaromatic drug treatment by BrdU incorporation. Untreated or nitroaromatic drug treated *T. brucei* was pulsed with 100 μM BrdU for 1 hour before parasites were harvested for flow cytometry analysis. In each condition (treated and untreated) cells were gated (A) and BrdU positive parasite populations were identified using an anti-BrdU antibody conjugated to Alexa488 alongside DAPI DNA content staining (B &C). The BrdU positive populations of nifurtimox (blue, 15μM), benznidazole (green, 180 μM), and fexinidazole (orange, 50 μM) treated (24 hours) parasites were then compared to untreated (UT, black) cells (D) and among drug treatment conditions by anti-BrdU histogram (E) and anti-BrdU vs. DAPI overlay (F).

### Analysis of DNA break formation during nitroaromatic drug treatments

When nitro drugs are activated by the trypanosome type I NTR enzyme, they are predicted to cause activation of ROS whose myriad of potential outcomes include DNA damage. Specifically, the oxidation of DNA bases that would then require repair to proceed with DNA synthesis, the completion of cell cycle, and cytokinesis(Chang et al., 2017). Mutated DNA bases can also result in the formation of DNA double-stranded breaks through replication fork collapse and other mechanisms (Truong et al., 2013; Iyer and Rhind, 2017). Because of the specific and nuanced differences that we have observed here between fexinidazole and related nitroaromatic trypanosome therapeutics, we sought to evaluate DNA break formation in a manner that was coordinated with cell cycle progression. The DNA histone H2A is present on DNA and is phosphorylated in the event of DNA damage to form γH2A-Phos, one of the most prominent and earliest markers of DNA damage in eukaryotes (Rogakou et al., 1998). Trypanosomes are highly divergent eukaryotes and, thus, have an altered phosphorylation site, H2A-Thr^130^, which has been well established in *T. brucei* as a marker for DNA damage by western blot and IF microscopy visualization of DNA break foci using a γH2A-Phos monoclonal antibody (Glover and Horn, 2012). To evaluate when DNA damage occurs during cell cycle, we sought to develop a γH2A-P-based flow cytometry assay using an anti-γH2A-P antibody conjugated to Alexa488 fluorophore that can be assayed along with DAPI staining to analyze the timing and extent of DNA break formation under different conditions. To establish a γH2A-P-based flow cytometry assay we compared UT unstained cells with γH2A-P-A488 stained parasites that were either UT or phleomycin (BLE) treated for 12 hours, a positive control for DNA break formation (Supplemental Figure 4). Populations of UT unstained, UT γH2A-P-A488 stained, and BLE treated γH2A-P-A488 stained populations were clearly separable on γH2A-P-A488 histograms, with 80% BLE treated cells shifting to a high γH2A-P+ staining population, compared with only 2% of UT (Supplemental Figure 4). Analysis of γH2A-P-A488 in comparison with DAPI allowed the specific identification of DNA break harboring parasites arising in both the G_1_ and G_2_ populations, relatively uniformly, upon BLE treatment. Overlay of unstained, γH2A (antibody stained), and γH2A-P+ (DNA break) populations graphed DAPI vs γH2A-P-A488 demonstrate the usefulness of this flow cytometry-based assay in the coordinated analysis of DNA break formation and cell cycle progression (Supplemental Figure 4).

DNA break formation was then analyzed during treatment with nifurtimox (15μM), benznidazole (180μM), and fexinidazole (50μM) at 12 hours (data not shown), 24 hours, and 48 hours post treatment. Little to no increase in DNA break formation was observed at 12 hours in all conditions and, thus, was not included in the associated data figure. Using the gating established based on BLE DNA damage induction (Supplemental Figure 4), we observed that nifurtimox and benznidazole resulted in a minor increase in γH2A-P+ (DNA damage) populations at 24 hours, 11% and 6% respectively (FIG. 7A). It is notable that when evaluated alongside DAPI staining, both nifurtimox and benznidazole result in a high staining G_1_ subpopulation of parasites containing DNA damage as early as 24 hours. By 48 hours the G_1_ DNA damaged subpopulation observed upon nifurtimox and benznidazole treatment is more pronounced and a new population DNA damaged parasites in S phase has emerged (FIG. 7B). By 48 hours, nifurtimox and benznidazole resulted in 26% and 27% DNA damaged parasites, respectively, across all cell cycle populations (FIG. 7B). Fexinidazole treatment also resulted in an increased populations of DNA damaged parasites, however the patterns of DNA break accumulation appears distinct from the related drugs analyzed (FIG. 8D). Specifically, at 24 hours, based on the gating we established, fexinidazole treatment resulted in a 20% increase in DNA break formation and 28% by 48 hours of treatment but without the specific G_1_ and S phase γH2A-P+ populations observed for nifurtimox and benznidazole (FIG. 7A & 7B). It is notable that while nifurtimox and benznidazole produce a clearly separable γH2A-P+ small peak after 24 hours, the larger γH2A-P+ population generated by fexinidazole is visualized more as a broadening of the γ-H_2_A+ population than a separate γH2A-P+ peak (FIG. 7A - γH2A-P histogram, 20% γH2A-P+). By 48 hours post fexinidazole treatment this peak has further broadened into a plateau accounting for 28% of parasites being γH2A-P+ (FIG. 7B). It is also notable that the γH2A-P peak arising from fexinidazole treatment (48 hours) also broadens in the negative direction (generally wider histogram than other samples), which might be associated with an increase in low staining AN parasites that occurred in this treatment condition (FIG. 7C). When the three drugs were compared, we observe that while all treatments result in an increase in DNA break formation, of similar proportions at 48 hours (26%, 27%, and 28%), the pattern of fexinidazole γH2A-P+ is distinct with respect to nifurtimox and benznidazole (FIG. 7D). Namely, nifurtimox and benznidazole have a specific accumulation of parasites in G_1_ harboring DNA breaks whereas fexinidazole has broader and more generally diffuse DNA break formation across G_1_ and G_2_, but not S phase (as the S phase population is reduced in this condition) (FIG. 7D). It is worth considering the possibility that DNA breaks in arising in G_1_ as a result of nifurtimox and benznidazole treatment may be the reason parasites do not proceed to G_2_ under these conditions (FIG. 3). The effects of fexinidazole treatment on DNA break formation must be considered in context with our observation that DNA synthesis is largely inhibited within 24 hours of the same treatment condition (FIG. 6), which will be considered further in the discussion (FIG. 8 – MODEL). In summary, while the anti-trypanosome nitroaromatic drugs tested result in a similar accumulation of DNA damage following 48 hours of treatment (26-28%), the cell cycle timing of DNA break formation in fexinidazole appears to be distinct from benznidazole and nifurtimox.

**FIGURE 7.**
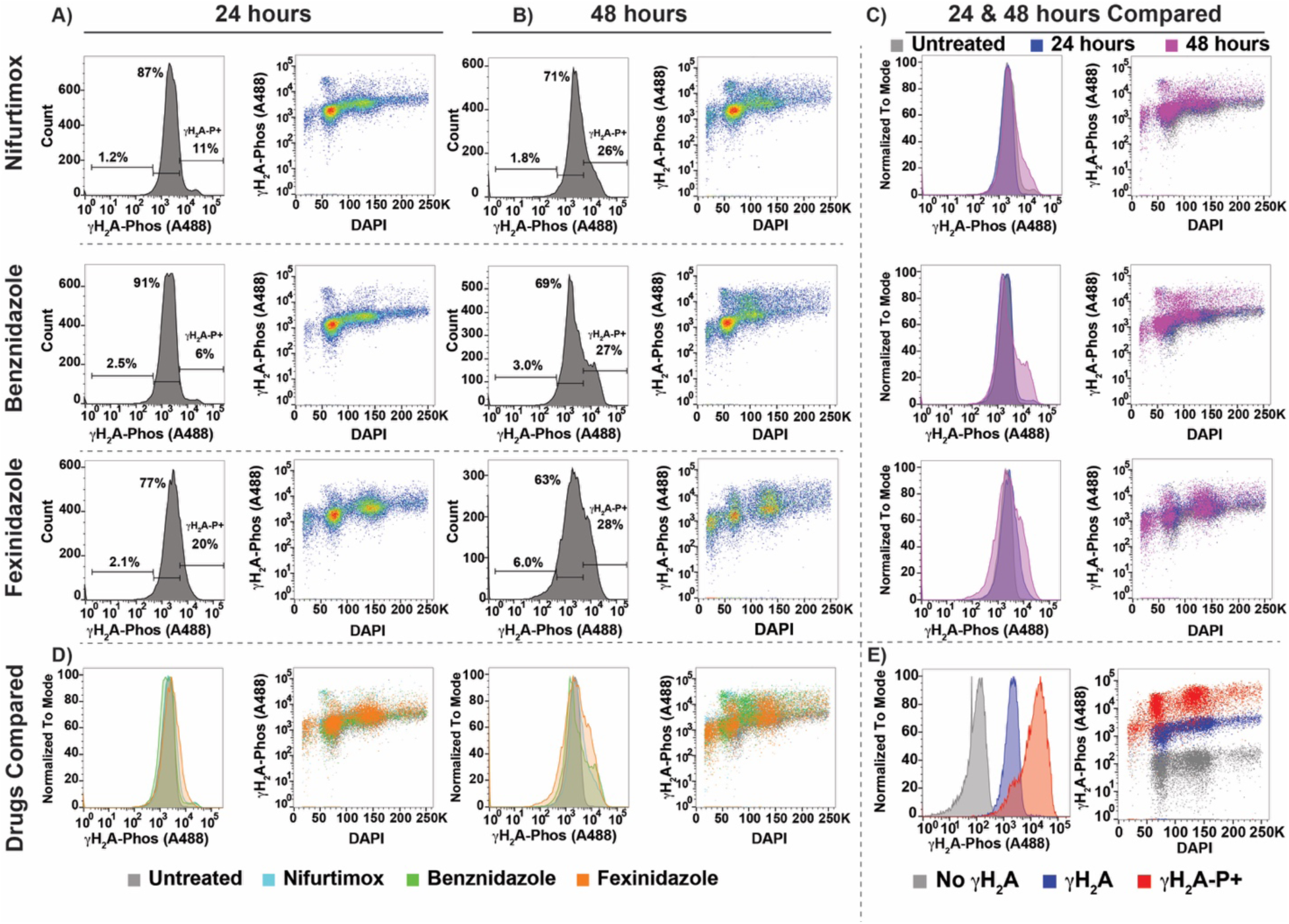
Evaluation of DNA damage during nitroaromatic drug treatment using. γ**H2A.** DNA damage was evaluated based on a *T. brucei* specific anti-γH2A-P (Alexa488 conjugated) antibody in conjunction with DAPI staining after treatment with nifurtimox (15 μM), benznidazole (180 μM), and fexinidazole (50 μM) at 24 and 48 hours (A & B). Flow cytometry data is displayed as anti-γH_2_A histograms or anti-γH_2_A vs. DAPI. C) The 24 and 48 hour time points are compared for each drug (24 hour blue and 48 hour purple) alongside untreated (anti-γH2A stained) parasites (grey). D) Nitroaromatic drugs (blue=nifurtimox, green=benznidazole, and orange=fexinidazole) are then compared at 24 and 48 hours. E) Gating and percent (%) of anti-γH2A-P positive cells were based on unstained (grey), anti-γH2A-P stained (navy), and a phleomycin positive DNA break control (red). For more information on anti-γH2A-P gating see Supplemental Figure 4.

**FIGURE 8.**
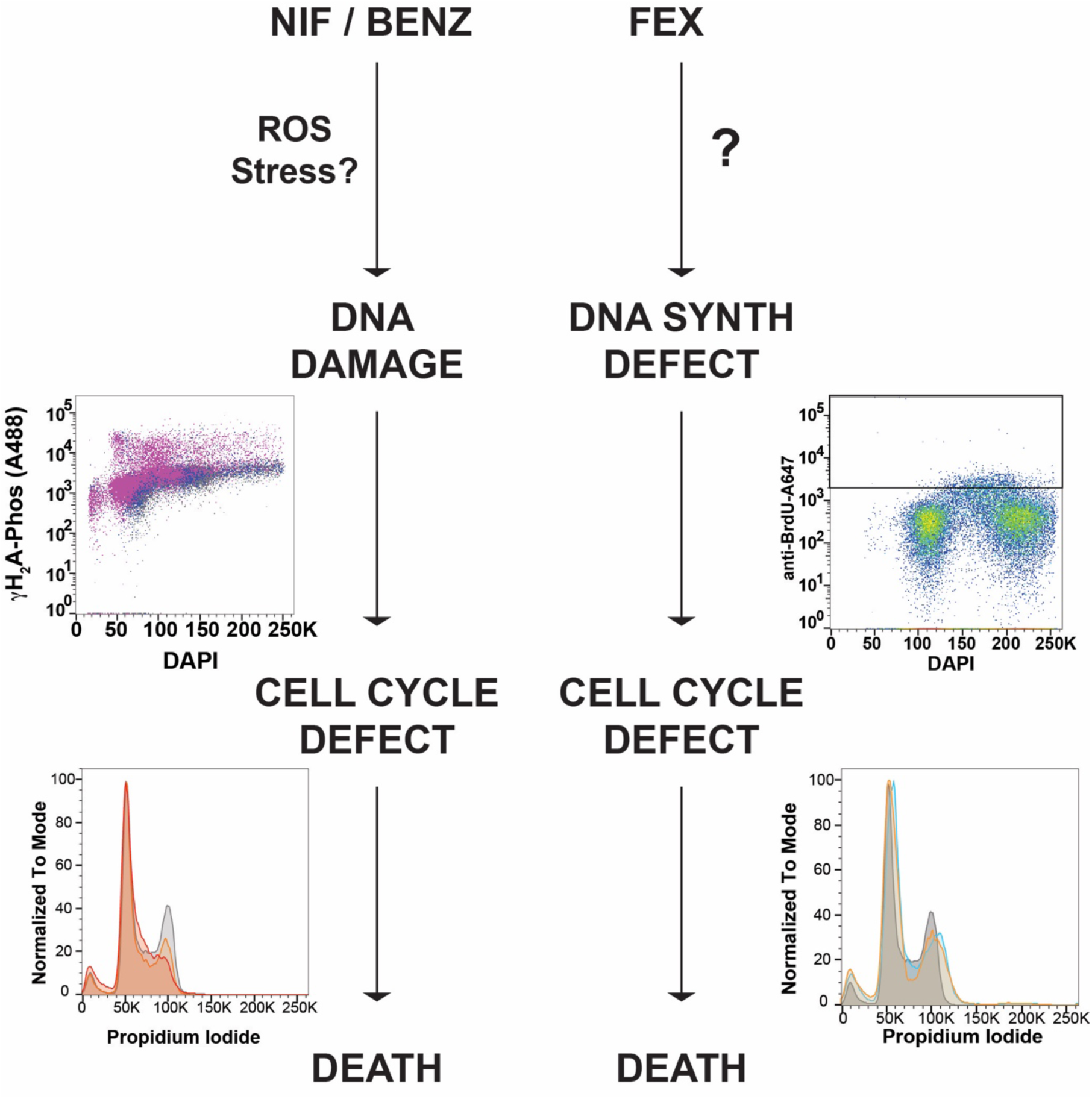
Model for the observed differences in nitroaromatic drug trypanocidal activity. Nifurtimox and benznidazole treatments progress similarly to form DNA damage, which appears to result in the loss of the G_2_ population, a resulting cell cycle stall, and death. In contrast, fexinidazole results in a distinct loss of DNA synthesizing cells, leading to loss of S phase cells, increased G_2_, and, ultimately, parasite death.

## 3. DISCUSSION

Fexinidazole is positioned to change the landscape of HAT treatment in sub-Saharan Africa by providing patients with an oral therapy to against African Sleeping Sickness for the first time. However, both current cornerstone therapies against second stage g-HAT, NECT and fexinidazole, now rely on nitroaromatic compounds leaving them vulnerable to drug resistance and cross-resistance (Sokolova et al., 2010b). Due to the efficacy of fexinidazole against apparently all kinetoplastida parasites, the drug is being actively considered as a possible treatment for American trypanosomiasis and multiple forms of Leishmania infections (Bahia et al., 2014; Patterson and Fairlamb, 2019; Mazzeti et al., 2021). While fexinidazole’s clinical approval may represent a considerable therapeutic advancement, there are major gaps in our understanding of how it kills trypanosomes and sources of drug resistance and nitroaromatic cross-resistance (Sokolova et al., 2010b; Hall et al., 2011; Wyllie et al., 2016b). This study sought to dissect specific cytotoxic outcomes arising from each of the clinically relevant nitroaromatic drugs: nifurtimox, benznidazole, and fexinidazole. In summary, we report that fexinidazole’s cytotoxicity is predominantly associated with defective DNA synthesis, whereas nifurtimox and benznidazole are more DNA damage associated (FIG. 8 – Model).

The potential for nitroaromatic compounds to result in DNA damage in trypanosomes has been reported since the 1980s for nifurtimox and benznidazole (Knox et al., 1981; Goijman and Stoppani, 1985). While these drugs can affect other macromolecules (RNA, proteins, and lipids) they have a particular propensity for damaging nucleotides. More recently, in the evaluation of eflornithine and nifurtimox metabolomics toward NECT development, nifurtimox resulted in increased concentrations of nucleotides and nucleobases, which was viewed as consistent with its hypothesized role of the unsaturated nitrile acting as a Michael acceptor (electrophile) (Vincent et al., 2012). In contrast, the same study showed that eflornithine predominantly effected the polyamine metabolism, thus demonstrating a lack of synergy in the molecular mechanism of each drug that supports their use combinatorially as NECT (Vincent et al., 2012). Despite decades of research and clinical usage of nifurtimox and benznidazole, their trypanocidal effects have not been thoroughly interrogated. Treatment of *T. brucei* in vitro with benznidazole recently demonstrated that RAD51 and MRE11 localize to the nucleus during drug treatment and that this timing was coordinated with an increase in γH2A-P by western blot analysis (Dattani et al., 2021), lending additional support to its predicted role in DNA damage formation.

The cell cycle data presented herein are the first to report that nifurtimox and benznidazole have similar cell cycle effects resulting in the loss of cells in G_2_ and an associated cytokinesis defect (as parasites do not progress to the 2K2N stage) (FIG. 3). Our observation that nifurtimox and benznidazole result in a specific, high γH_2_A-P+ signal, in G_1_ (FIG. 7) may suggest that these drugs initially cause DNA damage in G_1_ cells and that this hinders their inability to proceed to G_2_. This was distinct from fexinidazole, which displayed a general pattern of cells harboring DNA damage among both G_1_ and G_2_ populations (FIG. 7). These data suggest that nifurtimox and benznidazole treatment cause DNA damage in normally dividing cells (G_1_) that prevents them from completing cell cycle (G_2_ defect). Whereas fexinidazole’s cytotoxic effects may not arise as a result of DNA damage, but rather suffer DNA damage as a consequence of DNA synthesis inhibition (FIG. 8 – Model).

Within 24 hours of fexinidazole treatment we observed a nearly complete inhibition of DNA synthesis, as shown by BrdU incorporation assay (FIG. 6). Notably, inhibition of DNA synthesis has been suggested to be a powerful target of anti-trypanosome drugs (Otero et al., 2006). Previously, we reported that melarsoprol, an important drug in the treatment of second stage HAT, resulted in an early and potent effect on *T. brucei* DNA synthesis in vitro (Larson et al., 2021). In the same study we showed that while eflornithine and pentamidine do not have a negative effect on DNA synthesis, fexinidazole resulted in a considerable decrease in DNA synthesis (Larson et al., 2021), which support the results reported here. Melarsoprol is an arsenical drug that has a specific anti-trypanosome effect of binding trypanothione, the primary thiol carrier in these parasites (Fairlamb et al., 1989). Trypanothione is critical for many thiol-based redox reactions, including the reduction of ribonucleotide reductase to generate and maintain the dNTP pool required for DNA synthesis (Krauth-Siegel and Comini, 2008). Thus, we predicted that melarsoprol’s inhibition of DNA synthesis arose as a result of its inactivation of trypanothione depleting the dNTP pool. In fact, we were able to show that overexpression of the rate limiting step of trypanothione biosynthesis (γ-glutamyl cystine synthetase, GSH1) was able to recover DNA synthesis during melarsoprol treatment (Larson et al., 2021). This was not, however, the case for fexinidazole whose DNA synthesis defect was not altered upon GSH1 overexpression in the same study. Notably, melarsoprol treatment resulted in a decrease in the G_2_ population of parasites, which is not the case for fexinidazole, suggesting that DNA synthesis defects can result in alternative cell cycle outcomes. The specific details of their drug induced cell cycle defects may provide clues to the nature of their mechanisms of DNA synthesis inhibition.

The timing of events following drug treatment appears to be of considerable importance. It is likely that if nitroaromatic drugs were evaluated at later timepoints or in higher concentrations, any nuance between their cytotoxic effects would be challenging to observe. Recently, the cellular ultrastructure of *T. cruzi* treated with fexinidazole over a range of concentrations (similar to those used here) was evaluated after 72 hours resulting in detachment of the plasma membrane, unpacking of nuclear heterochromatin, and alterations in the kinetoplast and mitochondrion, among others (Zuma and Souza, 2022). Distinguishing between the characteristics of a dying cell and the cytotoxic events that lead to death can be challenging to disentangle. To address this challenge, we identified concentrations of drug that eliminate parasites in vitro by around day 5 of treatment and then evaluate their cytotoxic effects within 24 or 48 hours of treatment. While drug potency is still a factor, for instance nifurtimox appears more potent than benznidazole, and fexinidazole may be more potent than either, establishing a similar cell death and analysis timeline for each drug is our approach to mitigating this challenge. For instance, in Figure 1 we can observe that treatment in all 3 drugs at the highest concentrations parasites are able to grow in the first 24 hours. Within this time we can see a collection of phenotypes that align for nifurtimox and benznidazole, which are distinct from fexinidazole. Namely, nifurtimox and benznidazole begin to decrease the number of cells in G_2_, have populations of cell harboring DNA breaks in G_1_, and have little to no effect on DNA synthesis. Where, fexinidazole has little to no defect in the formation of G_2_ population, but a pronounced effect on DNA synthesis (FIG. 6), which is reflected in a decrease in populations of DNA synthesizing cells (FIG. 2 & 5). Presumably, at later stages of treatment, the outcomes of cell cycle stalling from various treatment conditions will devolve into various cell death outcomes, but this was not what was observed within the timing of the analysis herein.

While it does appear clear that fexinidazole’s MoA is linked to the DNA synthesis defect reported here, what is less clear is how and why fexinidazole chemistry results in the inhibition of DNA synthesis. Among the nitroaromatic drugs, nitro group itself is likely the most chemically reactive species. Prior to NTR bioactivation, nifurtimox has the nitro group in the 5 position of a 5-nitrofuran while benznidazole and fexinidazole are 2- and 5-nitroimidazoles, respectively. These chemical distinctions are likely less significant than their NTR activated forms, with nifurtimox generating open-chain nitriles, benznidazole generating reactive glyoxal (Patterson and Wyllie, 2014b), and fexinidazole’s NTR-activated products still unreported. It is notable that the cytotoxic effects of nifurtimox and benznidazole, reported herein, are remarkably similar when their NTR-based bioactivation is expected to generate significantly different chemical outcomes. An additional consideration for fexinidazole is that, as an oral therapy, it is further processed by host metabolism to generate fexinidazole sulfone and fexinidazole sulfoxide, whose distinct anti-trypanosome effects, if any, have not been thoroughly investigated. It is notable, that fexinidazole was invigorated as a therapy to answer the need for improvement over melarsoprol, which suffers from high host toxicity, yet both the new drug and the old display inhibition of DNA synthesis. Perhaps our findings support the proposal that inhibition of DNA synthesis is an ideal anti-trypanosome drug targeting strategy that can be further exploited. Because fexinidazole treatment appears prone to the generation of drug resistance, and nitro-drug cross resistance, perhaps these data could be applied to the generation of a combined drug treatment based on fexinidazole and a (preferably non-nitroaromatic) drug with alternative trypanocidal activity. Elucidating precisely how fexinidazole inhibits DNA synthesis and results in nitroaromatic drug cross-resistance will be the subject of future studies.

## 4. METHODS

### Culture and drug treatments of *T. brucei*

Bloodstream form *T. brucei* (*Lister427*) single marker (SM) parasites were used in all untreated and drug treated experiments herein (Wirtz et al., 1999), which were conducted in HMI-9 media in vitro (Hirumi and Hirumi, 1994). Supplier provided anti-trypanosome drugs, benznidazole (Sigma-Aldrich, 419656), eflornithine (United States Pharmacopeia, 1234249), fexinidazole (SelleckChem, S2600), and nifurtimox (Sigma-Aldrich, N3415), as powders that were resuspended in their designated solvent and stored at −20°C prior to use. For experimental use each drug was diluted in HMI-9 to their final concentration as shown for each experiment. Drug efficacy and parasite death during drug treatments were evaluated in three ways: 1) cumulative growth assays, 2) death flask assays, 3) EC_50_ determination by AlamarBlue (ThermoFisher, DAL1100) cell viability assay. 1) Cumulative growth assays were conducted in 12 well culture dishes in 2 mL of HMI-9 or HMI-9 with drug added. Parasites were seeded at 100,000 cell/mL, parasite growth recorded after 24 hours, and diluted back to 100,000 cell/mL every day for a period of 1 week. Recorded daily counts were then converted to total cumulative growth and graphed in GraphPad Prism. All experiments included at least three biological replicates and error bars, when shown on graphs, represent the mean and standard deviation based on at least 3 independent biological replicates. 2) “Death flask” assays were utilized to observe the timing of parasite death over 1-2 weeks as well as the occurrence of spontaneous drug resistance in under drug selection. Tissue culture flasks (T-25) were inoculated with 10,000 cells/mL in 5-10 mL of HMI-9 and counted daily by hemocytometry. Every 3 days, cultures were centrifuged (1,500 RPM, 10 minutes) to isolate parasites, and transferred to a new flask containing fresh media and drug in the appropriate drug treatment condition. Daily counts were recorded and graphed using GraphPad Prism. All experiments included at least three biological replicates and error bars, when shown on graphs, represent the mean and standard deviation based on at least 3 independent biological replicates. 3) EC_50_ determination was conducted in 96 well plate assays using AlamarBlue to determine cell viability during drug treatments. Parasites were plated at 25,000 cells / well and treated for 48 hours prior to the addition of 20 μL of Almarblue per well and 4 hours of incubation (manufacturers protocol optimized for *T. brucei*). Plates were then read on an SpectraMax i3x Multi-Mode Microplate Detection Platform at 590 nm emission and 530 nm excitation fluorescence wavelength. AlmarBlue florescence intensity was converted to percent (%) death using puromycin treated controls as 100% death based on the following formula: % dead = 1-(avg test abs – avg puro abs) / (avg UT abs– avg puro abs). The resulting data over a range of drug concentrations was then graphed to show % death vs. log of the nM concentration of drug using GraphPad Prism for at least 3 biological replicates. Note* average EC_50_ determination for benznidazole, fexinidazole, and nifurtimox shown in Supplemental Figure 1.

### Cell cycle analysis by flow cytometry and associated statistical analysis

Standard flow cytometry approaches were used to measure cell cycle progression using propidium iodide staining of approximately 5 million cells following formaldehyde fixation as described (Pozarowski and Darzynkiewicz, 2004). Cell cycle analysis, and all flow cytometry analyses performed herein, were performed using a BDFACSCelesta and FACSDiva software for acquisition analysis. The resulting data were then analyzed and visualized using the FlowJo analysis package. Cell cycle gating (AN, G_1_, S phase, G_2_, and multinucleated) generated in FlowJo for at least 3 biological replicates for each treatment condition then underwent statistical analysis using GraphPad Prism with multiple t-tests and unpaired with parametric test, pval<0.001 (***) or pval<0.01 (**).

### Immunofluorescence microscopy

*T. brucei* parasites were grown under shown conditions (UT or drug treated) in a manner that enabled the isolation of 5-10 million cells per condition. Parasites were isolated, resuspended in PBS and 1% formaldehyde and allowed to settle on poly-L-Lysine coated slides (Electron Microscopy Service, 63410-01). Parasites were then stained with an anti-VSG-2 monoclonal antibody (Hovel-Miner et al., 2012) conjugated to Alexa647 and mounted in Vectashield (Vector Laboratories, H-1800-10) containing 4′,6-diamidino-2-phenylindole (DAPI). Slides were visualized on a Leica DMi8 Inverted Fluorescent Microscope and images were analyzed using the Fiji (ImageJ) software package for the identification and counting of the number of nuclei (N) and kinetoplasts (K) per parasite for at least 100 parasite cells per condition. Final image preparation was completed in Adobe Photoshop in keeping with standard guidelines.

### DNA synthesis quantification by BrdU incorporation assay

Exponentially growing *T. brucei* was prepared for BrdU incorporation assays similar to previously described assays(Silva et al., 2017; Larson et al., 2021). Parasites were incubated with 100 μM of 5-bromo-2-Deoxyuridine (BrdU, Sigma Aldrich B5002) for 1 hour at 37°C, harvested by centrifugation, washed 2x in PBS, and fixed with 1% paraformaldehyde for 20 minutes. Fixed cells were then permeabilized with 0.1% Triton X-100 for 30 minutes, treated with 3 M HCl, washed 3 times in PBS, and incubated with anti-BrdU antibody conjugated with Alexa Fluor 647 (1:250 dilution, Thermo Fisher B35140), 1:1000 DAPI, and 0.5% BSA overnight. Prepared cells were then washed with PBS and resuspended for flow cytometry analysis as described.

### Flow cytometry analysis of DNA damage using anti-γ-H2A-Phos antibody

Anti-γH2A-Phos antisera were raised in rabbits using the KLH-conjugated phosphor-peptide, C-KHAKA[pT]PSV as described previously (Glover and Horn, 2012) using Thermo Fisher Scientific custom peptide synthesis and antibody services. The resulting anti-sera were purified and conjugated to Alexa488 using standard protocols. Validation of anti-γH2A-Phos detection of DNA damage in *T. brucei* was validated inhouse by western blot analysis and immunofluorescence microscopy of DNA break foci (not shown) prior to adaptation of the antibody to flow cytometry-based analysis of DNA damage herein. To evaluate DNA damage, as marked by γH2A histone phosphorylation, by flow cytometry *T. brucei* parasites were grown in the indicated UT or drug treated condition harvested by centrifugation and washed in cold PBS. Cells were then fixed in 1% paraformaldehyde and permeabilized in 0.1% Triton-X prior to staining with anti-γH2A-Phos-A488 used at a 1:100 dilution and DAPI. All samples were evaluated by flow cytometry to identify the cell gate, singlets, and the A488 populations associated with unstained, anti-γH2A-Phos-A488 stained, and DNA damaged (γH2A-P+) populations, positive control was phleomycin (BLE) treated for 12 hours (Supplemental Figure 4). Gating is either shown as an anti-γH2A-Phos-A488 histogram, used for % γH2A-P+ determination, or anti-γH2A-Phos-A488 vs. DAPI to visualize the formation of DNA damage over stages of cell cycle.

## ACKNOWLEDGMENTS

The members of the Hovel-Miner lab would like to acknowledge the excellent training and resources provided by the GWU SMHS Flow Cytometry Core, namely Dr. Gregory Cresswell, and the training resources and equipment provided by the GW Nanofabrication & Imaging Center, specifically Dr. Anastas Popratiloff who’s training was invaluable. Above all we would like to thank our funders who made this work possible through the following NIH NIAID funded research projects: R01AI170769 & R21AI174051.

## 5. SUPPLEMENTARY DATA AND SUPPLEMNTARY FIGURE LEGENDS

**SUP. FIG. 1.**
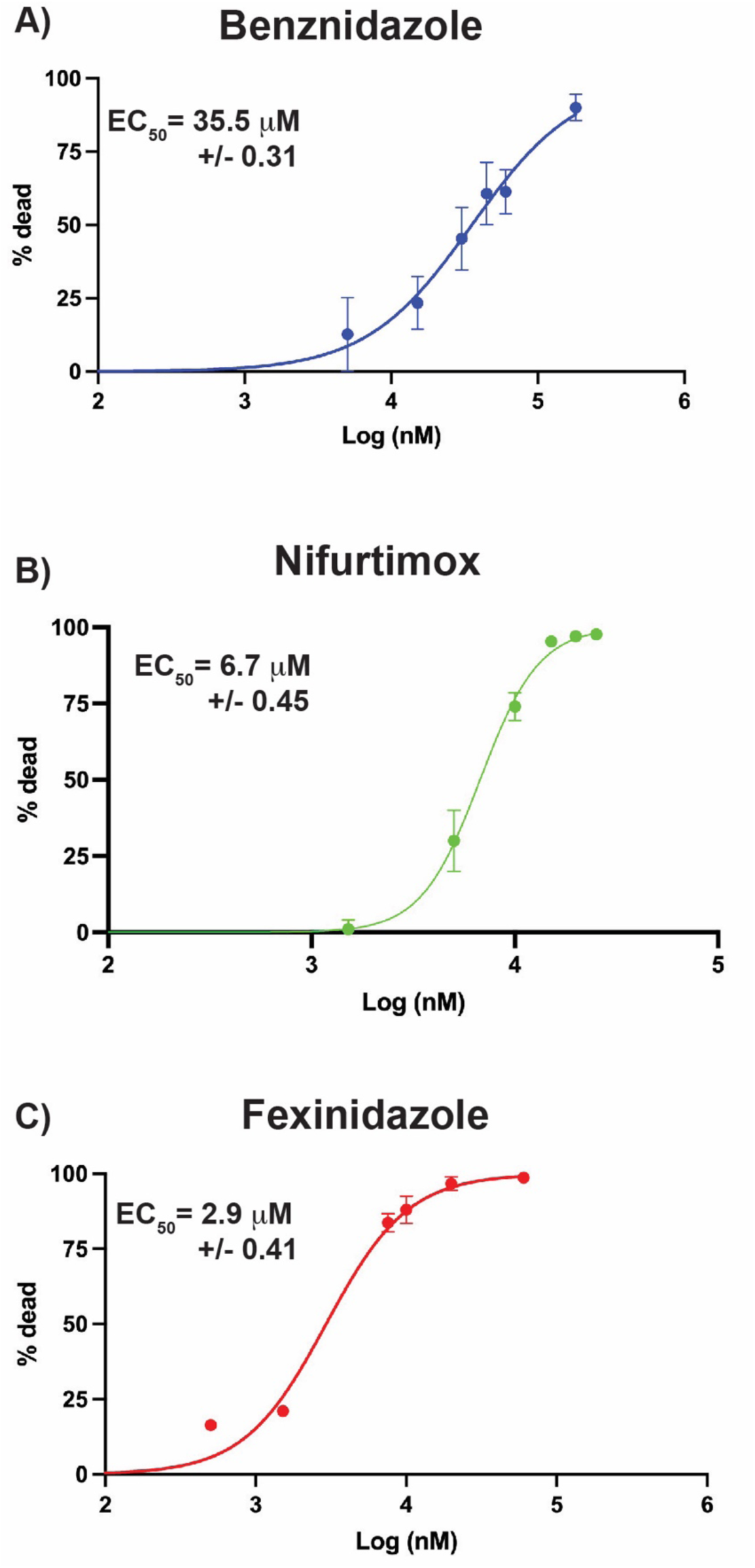
EC_50_ determination of anti-trypanosome nitroaromatic drugs. The EC_50_ for A) benznidazole (blue), B) nifurtimox (green), and C) fexinidazole (red) as determined during in vitro growth of *T. brucei* using AlamarBlue based cell viability assays. EC_50_ for each drug is shown inset on graph with standard deviation.

**SUP. FIG. 2.**
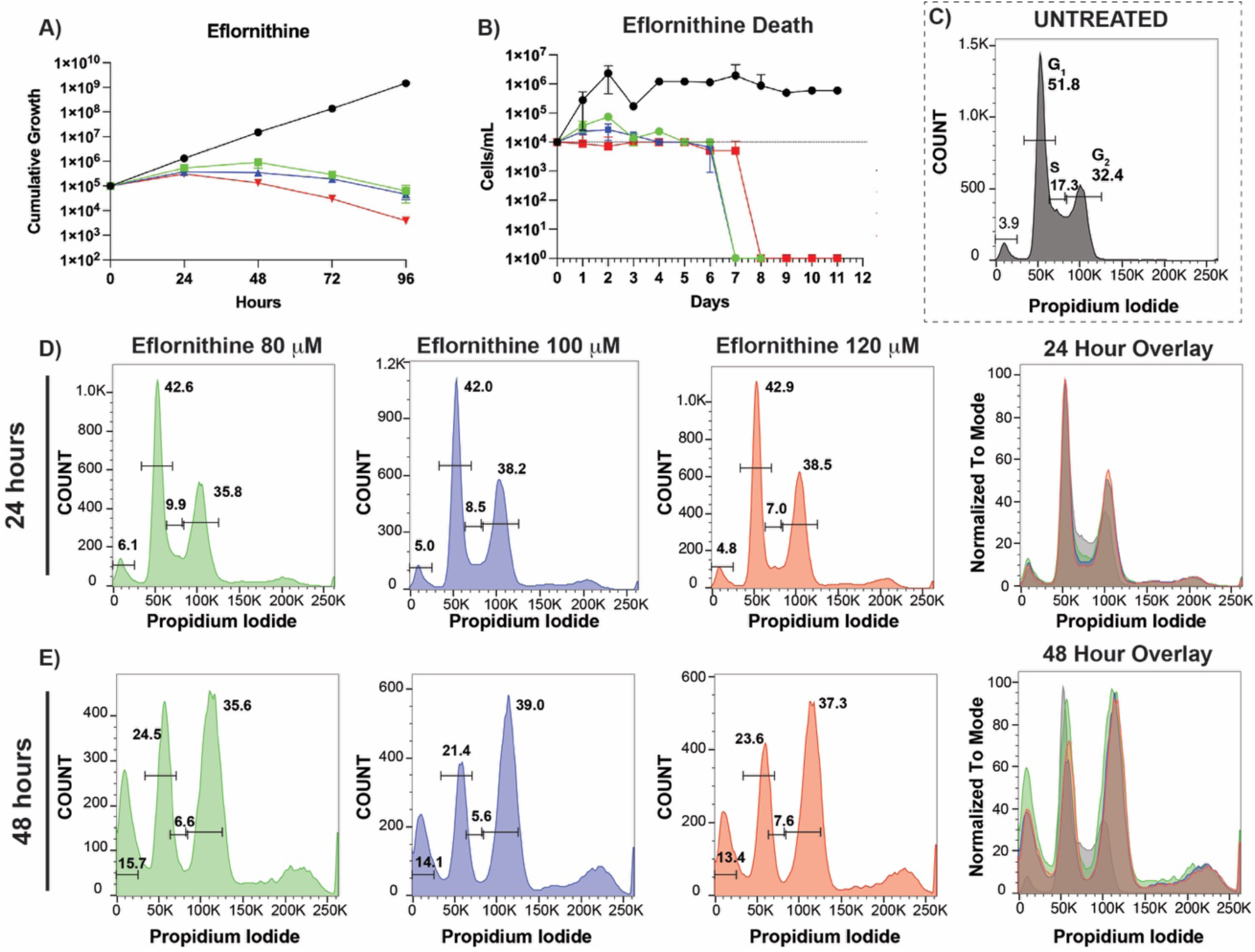
Eflornithine induced cytotoxicity. The effects of eflornithine were evaluated by A) cumulative growth assays and B) parasite death during treatment in flasks. Cell cycle was analyzed for C) Untreated parasites, after 24 hours (D) or 48 hours (E) of eflornithine treatment in low (green, 80 μM), mid (blue, 100 μM) and high (red, 120 μM) drug concentrations. Overlays with UT (grey) *T. brucei* shown to the right.

**SUP. FIG. 2.**
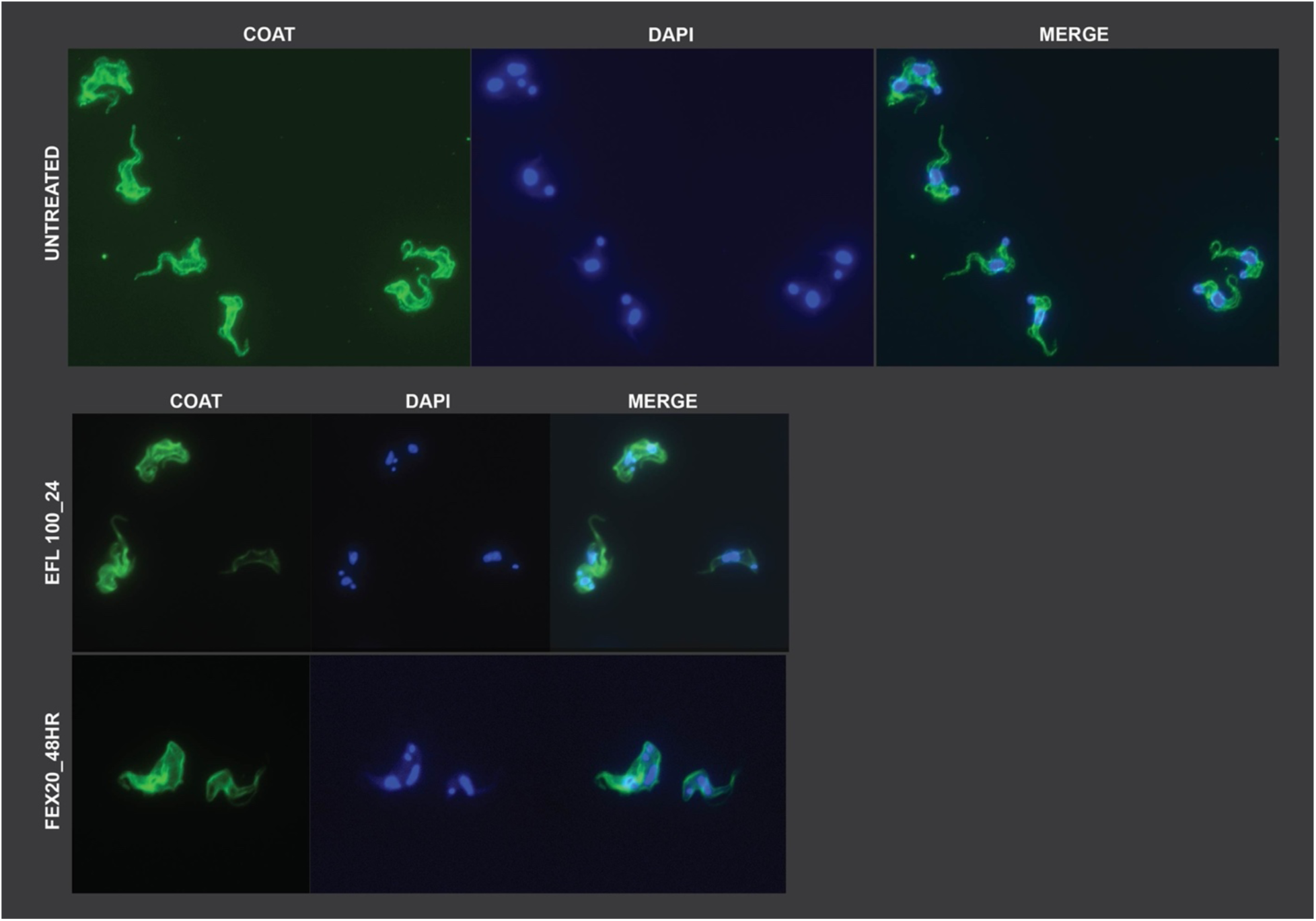
IF microscopy UT control and additional treatment conditions. IF microscopy of UT T. brucei is shown for anti-VSG2 (coat) antibody, DAPI, and merged image for one representative field. Representative images of drug induced cytotoxicity are also shown (bottom) for eflornithine (100 μM, 24 hours post-treatment) and fexinidazole (20 μM, 48 hours post-treatment).

**SUP. FIG. 4.**
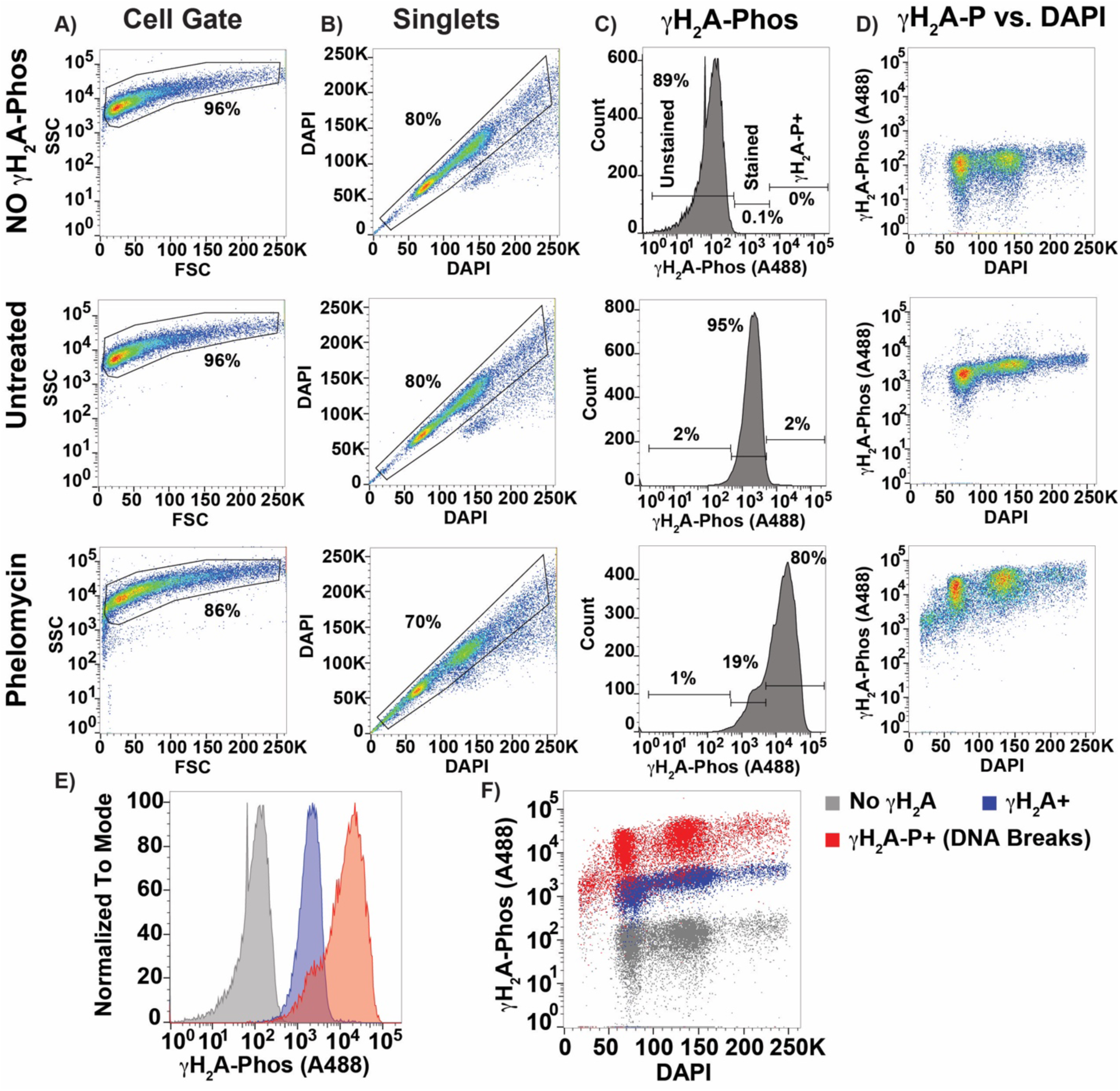
Flow cytometry gating of anti-γH2A-P analysis of DNA damage. (A) Cell gating, (B) DAPI-based doublet discrimination, (C) γH2A-Alexa488 histograms, and (D) γH2A-Alexa488 vs. DAPI are shown for unstained cells (NO γH2A), untreated γH2A-Alexa488 stained cells, and phelomycin (BLE) γH2A-Alexa488 treated cells, as a positive control for DNA damage. (E) Histograms of No γH2A (grey), γH2A+ untreated (blue), and γH2A-P+ (red, DNA Breaks) were compared to established gating for DNA damaged populations. (F) No γH2A (grey), γH2A+ untreated (blue), and γH2A-P+ (red, DNA Breaks) are shown vs. DAPI to illustrate the formation of DNA breaks during cell cycle.

## Notes

### Competing Interest Statement

The authors have declared no competing interest.

## 6. REFERENCES

Alirol, E., Schrumpf, D., Heradi, J. A., Riedel, A., Patoul, C. de, Quere, M., et al. (2013). Nifurtimox-Eflornithine Combination Therapy for Second-Stage Gambiense Human African Trypanosomiasis: Médecins Sans Frontières Experience in the Democratic Republic of the Congo. Clin Infect Dis 56, 195–203. doi: 10.1093/cid/cis886.

Babokhov, P., Sanyaolu, A. O., Oyibo, W. A., Fagbenro-Beyioku, A. F., and Iriemenam, N. C. (2013). A current analysis of chemotherapy strategies for the treatment of human African trypanosomiasis. Pathog Glob Health 107, 242–252. doi: 10.1179/2047773213y.0000000105.

Bahia, M. T., Nascimento, A. F. S., Mazzeti, A. L., Marques, L. F., Gonçalves, K. R., Mota, L. W. R., et al. (2014). Antitrypanosomal Activity of Fexinidazole Metabolites, Potential New Drug Candidates for Chagas Disease. Antimicrob. Agents Chemother. 58, 4362–4370. doi: 10.1128/aac.02754-13.

Chang, H. H. Y., Pannunzio, N. R., Adachi, N., and Lieber, M. R. (2017). Non-homologous DNA end joining and alternative pathways to double-strand break repair. Nat Rev Mol Cell Bio 18, 495–506. doi: 10.1038/nrm.2017.48.

Dattani, A., Drammeh, I., Mahmood, A., Rahman, M., Szular, J., and Wilkinson, S. R. (2021). Unraveling the antitrypanosomal mechanism of benznidazole and related 2-nitroimidazoles: From prodrug activation to DNA damage. Mol Microbiol 116, 674–689. doi: 10.1111/mmi.14763.

Deeks, E. D. (2019a). Fexinidazole: First Global Approval. Drugs 79, 215–220. doi: 10.1007/s40265-019-1051-6.

Deeks, E. D. (2019b). Fexinidazole: First Global Approval. Drugs 79, 215–220. doi: 10.1007/s40265-019-1051-6.

Deeks, E. D., and Lyseng-Williamson, K. A. (2019). Fexinidazole in human African trypanosomiasis: a profile of its use. Drugs Ther Perspect 35, 529–535. doi: 10.1007/s40267-019-00672-2.

Fairlamb, A. H., Henderson, G. B., and Cerami, A. (1989). Trypanothione is the primary target for arsenical drugs against African trypanosomes. Proc National Acad Sci 86, 2607–2611. doi: 10.1073/pnas.86.8.2607.

Glover, L., and Horn, D. (2012). Trypanosomal histone γH2A and the DNA damage response. Mol Biochem Parasit 183–341, 78–83. doi: 10.1016/j.molbiopara.2012.01.008.

Goijman, S. G., and Stoppani, A. O. M. (1985). Effects of nitroheterocyclic drugs on macromolecule synthesis and degradation in Trypanosoma cruzi. Biochem. Pharmacol. 34, 1331–1336. doi: 10.1016/0006-2952(85)90514-3.

Hall, B. S., Bot, C., and Wilkinson, S. R. (2011). Nifurtimox Activation by Trypanosomal Type I Nitroreductases Generates Cytotoxic Nitrile Metabolites*. J. Biol. Chem. 286, 13088–13095. doi: 10.1074/jbc.m111.230847.

Hall, B. S., and Wilkinson, S. R. (2011). Activation of benznidazole by trypanosomal type I nitroreductases results in glyoxal formation. Antimicrob. agents Chemother. 56, 115–23. doi: 10.1128/aac.05135-11.

Hammarton, T. C. (2007). Cell cycle regulation in Trypanosoma brucei. Mol Biochem Parasit 153, 1–8. doi: 10.1016/j.molbiopara.2007.01.017.

Hirumi, H., and Hirumi, K. (1994). Axenic culture of African trypanosome bloodstream forms. Parasitol Today 10, 80–84. doi: 10.1016/0169-4758(94)90402-2.

Hovel-Miner, G. A., Boothroyd, C. E., Mugnier, M., Dreesen, O., Cross, G. A. M., and Papavasiliou, F. N. (2012). Telomere Length Affects the Frequency and Mechanism of Antigenic Variation in Trypanosoma brucei. Plos Pathog 8, e1002900. doi: 10.1371/journal.ppat.1002900.

Iyer, D. R., and Rhind, N. (2017). Replication fork slowing and stalling are distinct, checkpoint-independent consequences of replicating damaged DNA. Plos Genet 13, e1006958. doi: 10.1371/journal.pgen.1006958.

Jones, N. G., Thomas, E. B., Brown, E., Dickens, N. J., Hammarton, T. C., and Mottram, J. C. (2014). Regulators of Trypanosoma brucei cell cycle progression and differentiation identified using a kinome-wide RNAi screen. Plos Pathog 10, e1003886. doi: 10.1371/journal.ppat.1003886.

Knox, R. J., Knight, R. C., and Edwards, D. I. (1981). Interaction of nitroimidazole drugs with DNA in vitro: Structure-activity relationships. Br. J. Cancer 44, 741–745. doi: 10.1038/bjc.1981.261.

Krauth-Siegel, R. L., and Comini, M. A. (2008). Redox control in trypanosomatids, parasitic protozoa with trypanothione-based thiol metabolism. Biochim Biophys Acta 1780, 1236–48. doi: 10.1016/j.bbagen.2008.03.006.

Larson, S., Carter, M., and Hovel-Miner, G. (2021). Effects of trypanocidal drugs on DNA synthesis: new insights into melarsoprol growth inhibition. Parasitology 148, 1143–1150. doi: 10.1017/s0031182021000317.

Mazzeti, A. L., Gonçalves, K. R., Mota, S. L. A., Pereira, D. E., Diniz, L. de F., and Bahia, M. T. (2021). Combination therapy using nitro compounds improves the efficacy of experimental Chagas disease treatment. Parasitology 148, 1320–1327. doi: 10.1017/s0031182021001001.

Otero, L., Vieites, M., Boiani, L., Denicola, A., Rigol, C., Opazo, L., et al. (2006). Novel Antitrypanosomal Agents Based on Palladium Nitrofurylthiosemicarbazone Complexes: DNA and Redox Metabolism as Potential Therapeutic Targets †. J. Med. Chem. 49, 3322–3331. doi: 10.1021/jm0512241.

Patterson, S., and Fairlamb, A. H. (2019). Current and Future Prospects of Nitro-compounds as Drugs for Trypanosomiasis and Leishmaniasis. Curr. Med. Chem. 26, 4454–4475. doi: 10.2174/0929867325666180426164352.

Patterson, S., and Wyllie, S. (2014a). Nitro drugs for the treatment of trypanosomatid diseases: past, present, and future prospects. Trends Parasitol 30, 289–298. doi: 10.1016/j.pt.2014.04.003.

Patterson, S., and Wyllie, S. (2014b). Nitro drugs for the treatment of trypanosomatid diseases: past, present, and future prospects. Trends Parasitol. 30, 289–298. doi: 10.1016/j.pt.2014.04.003.

Pozarowski, P., and Darzynkiewicz, Z. (2004). “Analysis of Cell Cycle by Flow Cytometry,” in Methods in molecular biology (Clifton, N.J.)., ed. [“Axel H. Schönthal”] (Totowa, NJ: Humana Press), 301– 311. doi: 10.1385/1-59259-811-0:301.

Raether, W., and Seidenath, H. (1983). The activity of fexinidazole (HOE 239) against experimental infections with Trypanosoma cruzi, trichomonads and Entamoeba histolytica. Ann. Trop. Med. Parasitol. 77, 13–26. doi: 10.1080/00034983.1983.11811668.

Rogakou, E. P., Pilch, D. R., Orr, A. H., Ivanova, V. S., and Bonner, W. M. (1998). DNA Double-stranded Breaks Induce Histone H2AX Phosphorylation on Serine 139*. J. Biol. Chem. 273, 5858– 5868. doi: 10.1074/jbc.273.10.5858.

Rycker, M. D., Wyllie, S., Horn, D., Read, K. D., and Gilbert, I. H. (2023). Anti-trypanosomatid drug discovery: progress and challenges. Nat. Rev. Microbiol. 21, 35–50. doi: 10.1038/s41579-022-00777-y.

Shahi, S. K., Krauth-Siegel, R. L., and Clayton, C. E. (2002). Overexpression of the putative thiol conjugate transporter TbMRPA causes melarsoprol resistance in Trypanosoma brucei. Mol Microbiol 43, 1129–1138. doi: 10.1046/j.1365-2958.2002.02831.x.

Silva, M. S., Muñoz, P. A. M., Armelin, H. A., and Elias, M. C. (2017). Differences in the Detection of BrdU/EdU Incorporation Assays Alter the Calculation for G1, S, and G2 Phases of the Cell Cycle in Trypanosomatids. J Eukaryot Microbiol 64, 756–770. doi: 10.1111/jeu.12408.

Sokolova, A. Y., Wyllie, S., Patterson, S., Oza, S. L., Read, K. D., and Fairlamb, A. H. (2010a). Cross-Resistance to Nitro Drugs and Implications for Treatment of Human African Trypanosomiasis. Antimicrob Agents Ch 54, 2893–2900. doi: 10.1128/aac.00332-10.

Sokolova, A. Y., Wyllie, S., Patterson, S., Oza, S. L., Read, K. D., and Fairlamb, A. H. (2010b). Cross-Resistance to Nitro Drugs and Implications for Treatment of Human African Trypanosomiasis. Antimicrob. Agents Chemother. 54, 2893–2900. doi: 10.1128/aac.00332-10.

Truong, L. N., Li, Y., Shi, L. Z., Hwang, P. Y.-H., He, J., Wang, H., et al. (2013). Microhomology-mediated End Joining and Homologous Recombination share the initial end resection step to repair DNA double-strand breaks in mammalian cells. Proc National Acad Sci 110, 7720–7725. doi: 10.1073/pnas.1213431110.

Vincent, I. M., Creek, D. J., Burgess, K., Woods, D. J., Burchmore, R. J. S., and Barrett, M. P. (2012). Untargeted Metabolomics Reveals a Lack Of Synergy between Nifurtimox and Eflornithine against Trypanosoma brucei. PLoS Neglected Trop. Dis. 6, e1618. doi: 10.1371/journal.pntd.0001618.

Whitmore, G. F., and Varghese, A. J. (1986). The biological properties of reduced nitroheterocyclics and possible underlying biochemical mechanisms. Biochem. Pharmacol. 35, 97–103. doi: 10.1016/0006-2952(86)90565-4.

Wilkinson, S. R., Taylor, M. C., Horn, D., Kelly, J. M., and Cheeseman, I. (2008). A mechanism for cross-resistance to nifurtimox and benznidazole in trypanosomes. Proc. Natl. Acad. Sci. 105, 5022– 5027. doi: 10.1073/pnas.0711014105.

Wirtz, E., Leal, S., Ochatt, C., and Cross, G. M. (1999). A tightly regulated inducible expression system for conditional gene knock-outs and dominant-negative genetics in Trypanosoma brucei. Mol Biochem Parasit 99, 89–101. doi: 10.1016/s0166-6851(99)00002-x.

Wyllie, S., Foth, B. J., Kelner, A., Sokolova, A. Y., Berriman, M., and Fairlamb, A. H. (2016a). Nitroheterocyclic drug resistance mechanisms in Trypanosoma brucei. J Antimicrob Chemoth 71, 625–634. doi: 10.1093/jac/dkv376.

Wyllie, S., Foth, B. J., Kelner, A., Sokolova, A. Y., Berriman, M., and Fairlamb, A. H. (2016b). Nitroheterocyclic drug resistance mechanisms in Trypanosoma brucei. J. Antimicrob. Chemother. 71, 625–634. doi: 10.1093/jac/dkv376.

Zuma, A. A., and Souza, W. de (2022). Fexinidazole interferes with the growth and structural organization of Trypanosoma cruzi. Sci. Rep. 12, 20388. doi: 10.1038/s41598-022-23941-z.

Zuma, N. H., Aucamp, J., and N’Da, D. D. (2019). An update on derivatisation and repurposing of clinical nitrofuran drugs. Eur. J. Pharm. Sci. 140, 105092. doi: 10.1016/j.ejps.2019.105092.

