## Supplementary figures and images for "Fexinidazole induced cytotoxicity is distinct from related anti-trypanosome nitroaromatic drugs"

### Supplemental Figure 1

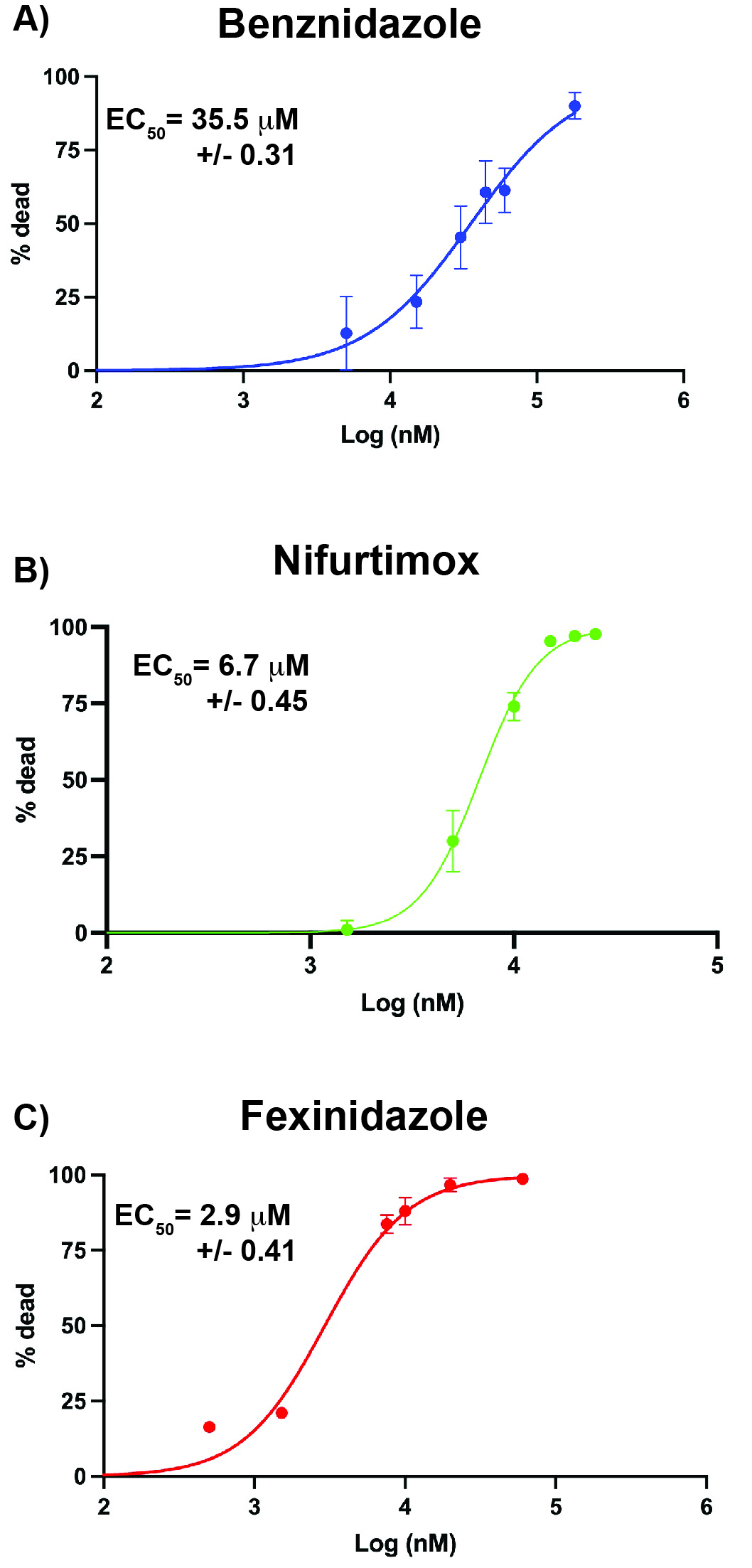

### Supplemental Figure 2

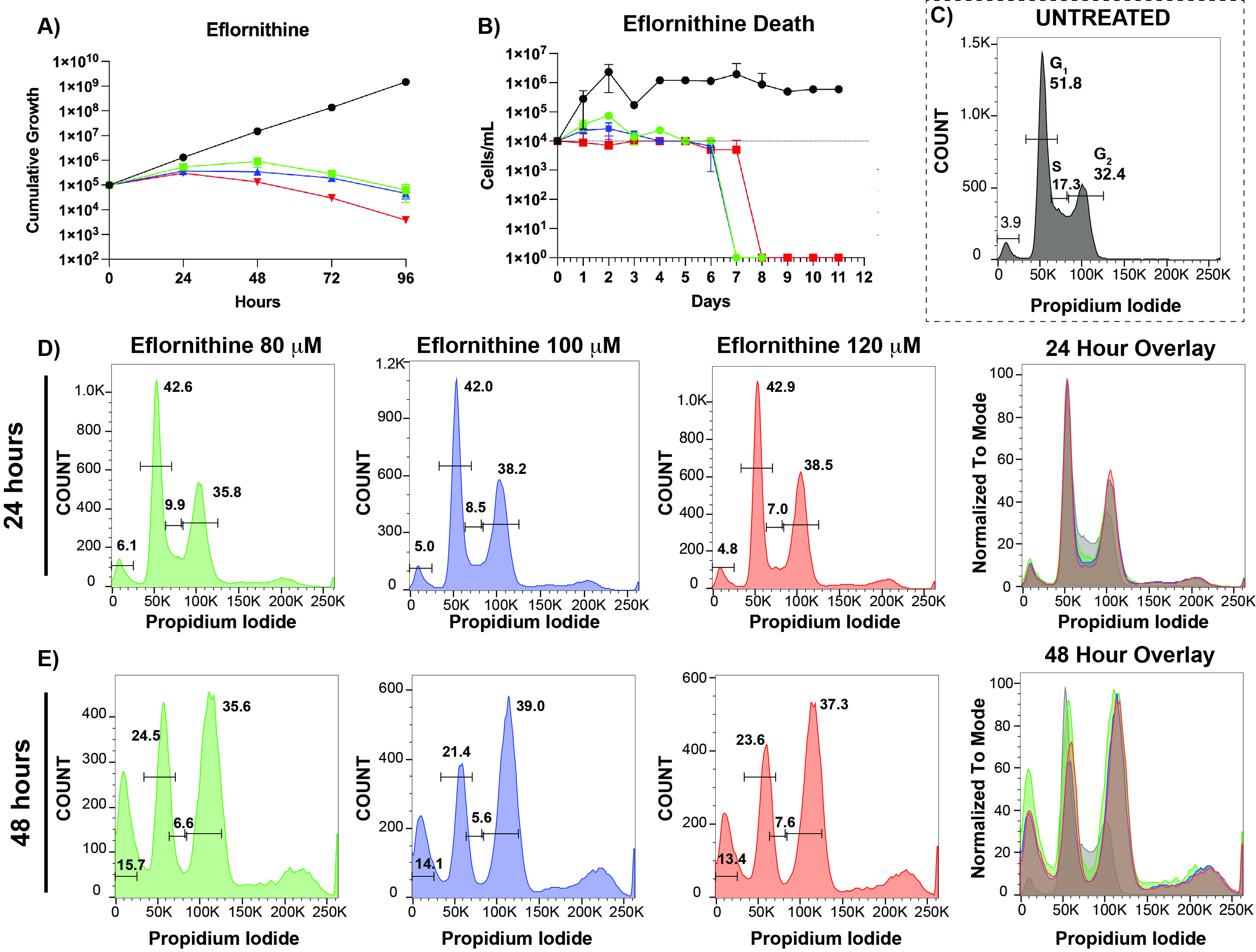

### Supplemental Figure 3

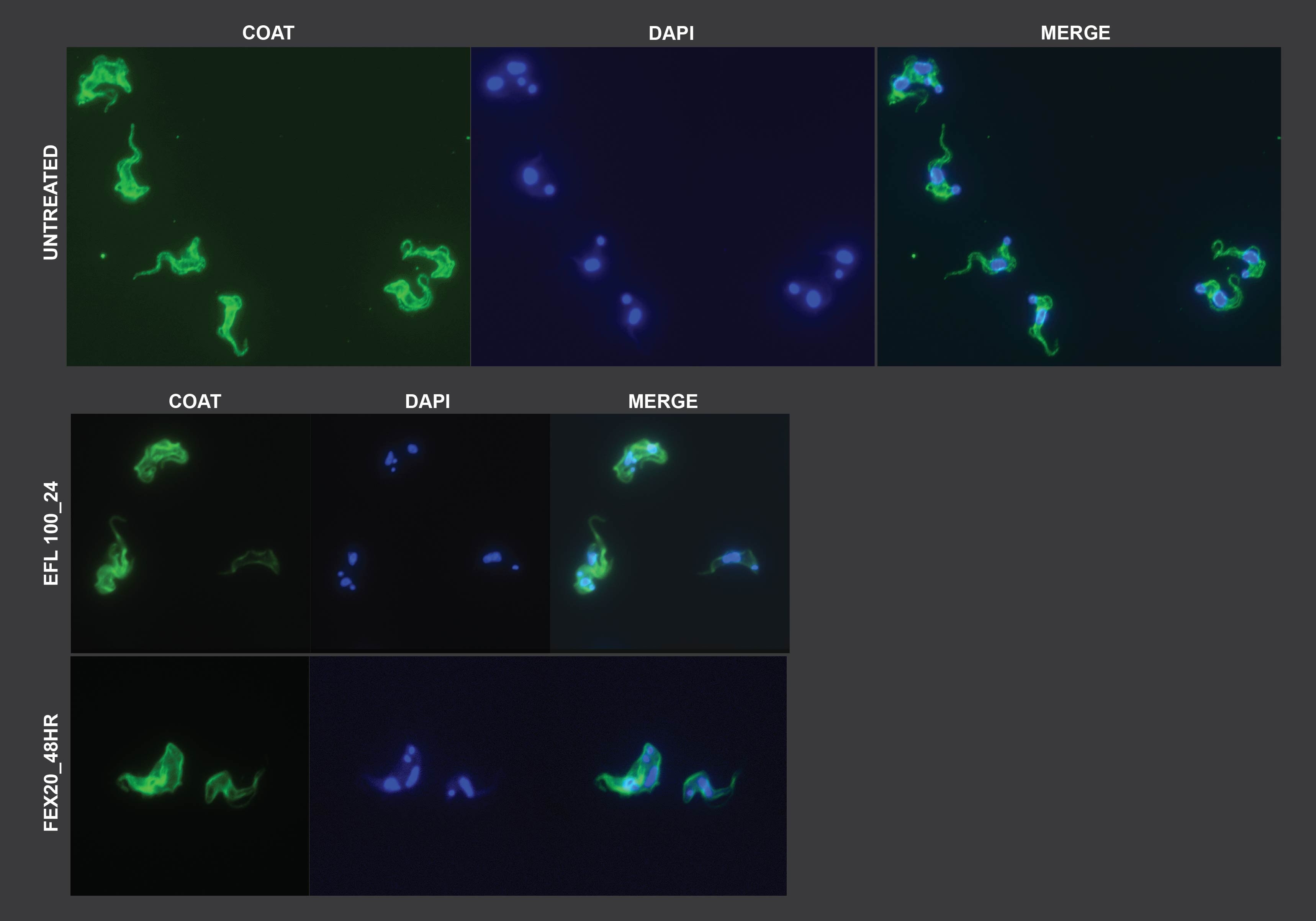

### Supplemental Figure 4

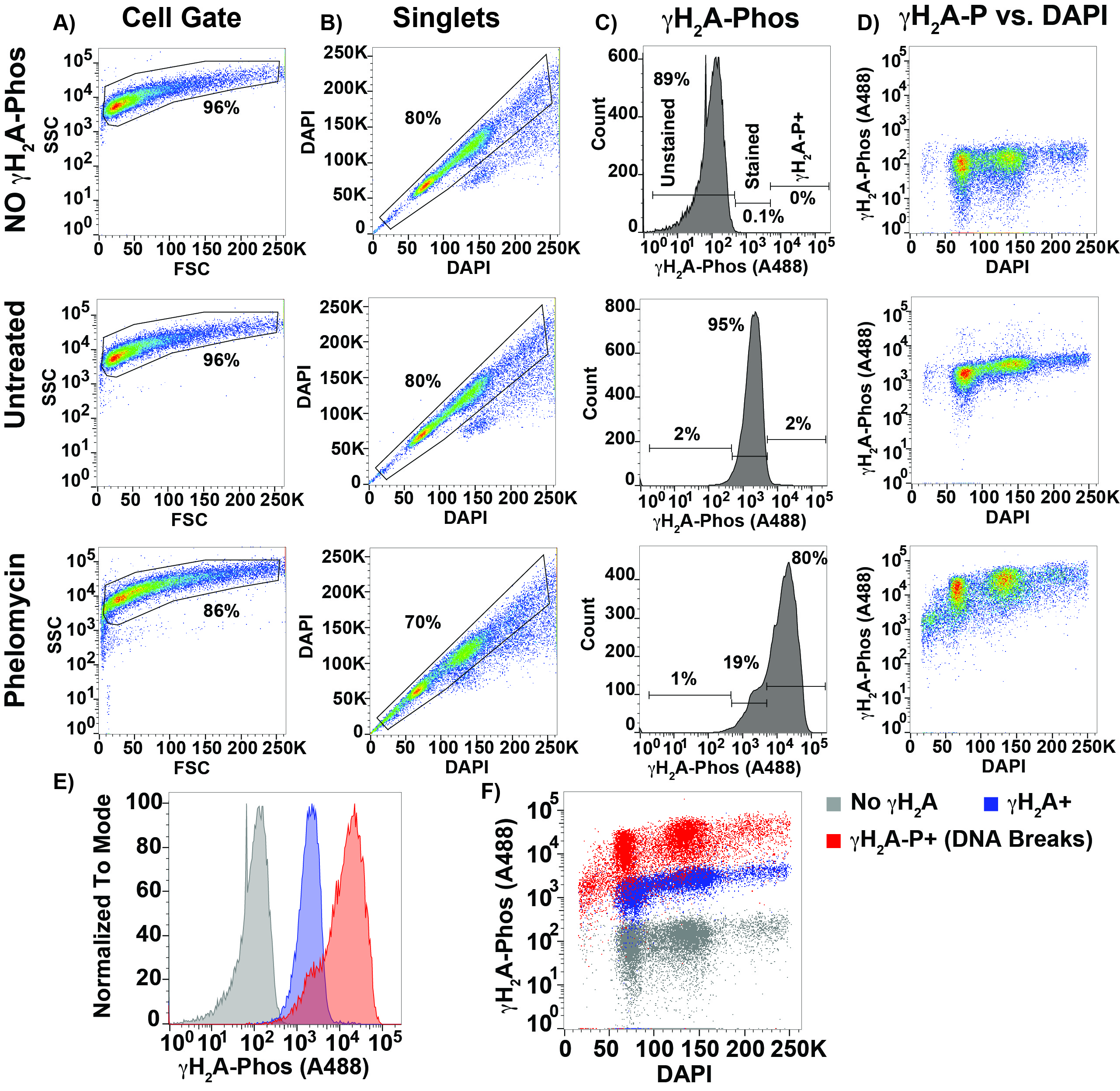
